# Life cycle-coupled evolution of mitosis in close relatives of animals

**DOI:** 10.1101/2023.05.10.540163

**Authors:** Hiral Shah, Marine Olivetta, Chandni Bhickta, Paolo Ronchi, Monika Trupinić, Eelco C. Tromer, Iva M. Tolić, Yannick Schwab, Omaya Dudin, Gautam Dey

## Abstract

Eukaryotes have evolved towards one of two extremes along a spectrum of strategies for remodelling the nuclear envelope (NE) during cell division: disassembling the NE in an open mitosis or constructing an intranuclear spindle in a closed mitosis. Both classes of mitotic remodelling involve key differences in the core division machinery, but the evolutionary reasons for adopting a specific mechanism are unclear. Here, we use an integrated comparative genomics and ultrastructural imaging approach to investigate mitotic strategies in Ichthyosporea, close relatives of animals and fungi. We show that species within this clade have diverged towards either a fungal-like closed or an animal-like open mitosis, most likely to support distinct multi- or uninucleated states. Our results suggest that multinucleated life cycles favour the evolution of closed mitosis.

**One-Sentence Summary:** Mitotic specialization in animal relatives reveal that multinucleated life cycles favor the evolution of closed mitosis

## Main Text

Eukaryotic mitosis relies on a tight coordination between chromosome segregation and the remodelling of the nuclear compartment to ensure the fidelity of nuclear division and genome inheritance (*1, 2*). Two classes of nuclear remodelling have been widely investigated: open mitosis (*3*), in which the nuclear envelope (NE) is disassembled at mitotic entry and reassembled following chromosome segregation, and closed mitosis (*4–7*), in which the nuclear compartment retains its identity throughout division (Fig. 1A). Although open and closed mitosis have each likely evolved independently multiple times in different branches of the eukaryotic tree (*8, 9*), with many unique lineage-specific adaptations resulting in a broad distribution of intermediates from fully open to fully closed (*1, 10*), the reasons that species tend towards one or the other mitotic strategy are not well understood. Studies, primarily in mammalian and yeast models, suggest that open and closed mitosis require distinct adaptations in key structural components of the division machinery (*10*), including the microtubule organising centre (MTOC) (*11*), the spindle (*12, 13*), the NE (*14, 15*) and the kinetochore (*16–18*). For example, building an intranuclear spindle in a closed mitosis must be accompanied by NE fenestration to allow insertion of the MTOC (*19*). On the other hand, open mitosis requires distinct interphase and post-mitotic mechanisms for the insertion of new nuclear pore complexes (NPCs) into the NE (*20*). These significant differences in the core division machinery imply that biophysical constraints linked to the mode of mitosis are encoded in the genome, an observation we can leverage through comparative genomics to provide testable predictions for mitotic phenotypes. Experimentally verifying these predicted phenotypes in a broad range of non-model species allows us to ask whether constraints imposed by ecological niche and life cycle could drive species towards either open or closed mitosis.

**Figure 1.**
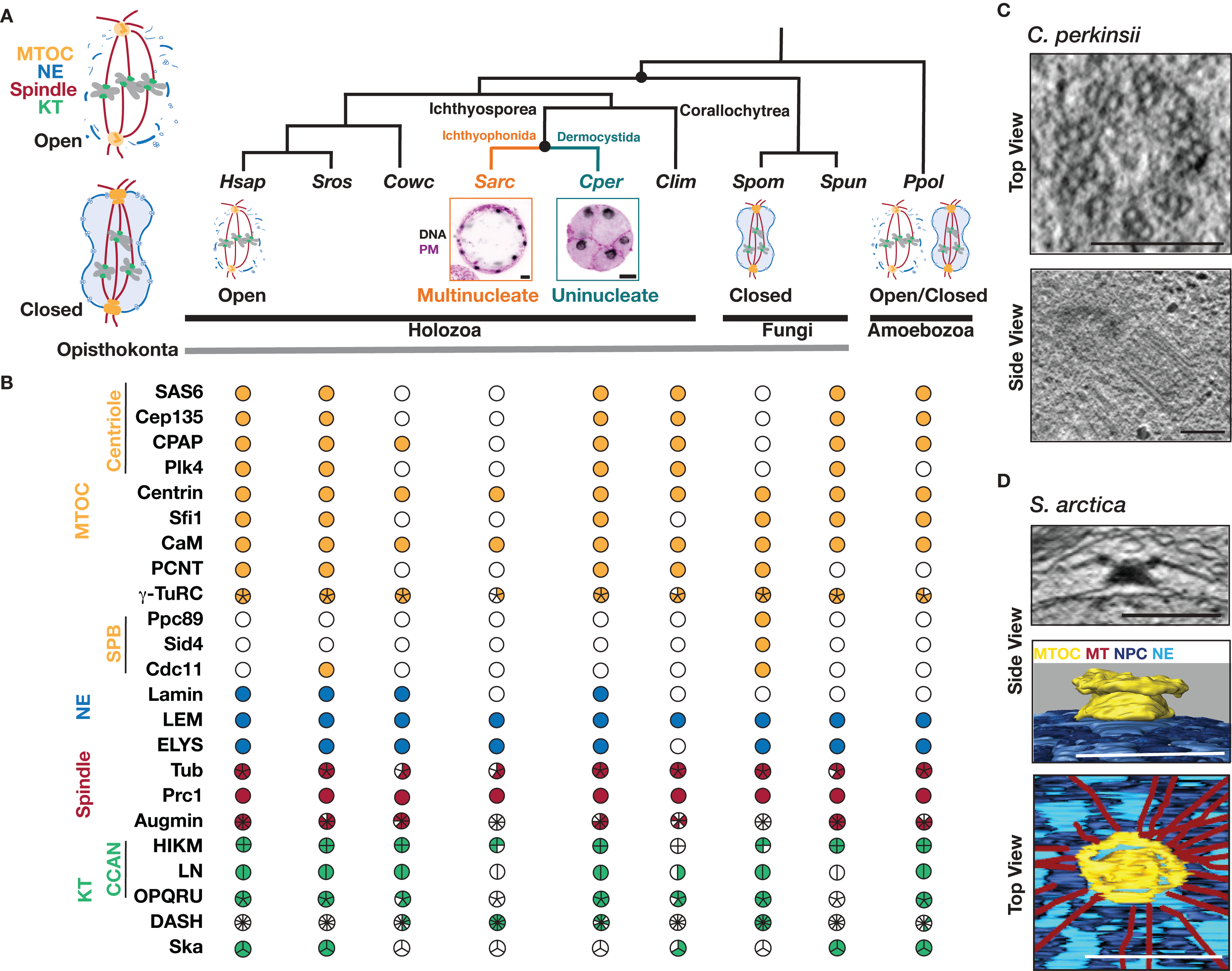
Divergence of mitotic machinery within Ichthyosporea. (**A**) Cladogram of opisthokonts, highlighting the position of Ichthyosporea between well-studied animal and fungal model systems. Schematics depicting the mitotic strategies, open or closed, of the represented opisthokont and amoebozoan species. Maximum intensity projections show differences in life cycles and the uni- and multinucleated states of *C. perkinsii* (*Cper*) and *S. arctica* (*Sarc*), respectively. Cells are labelled for plasma membrane (magenta) and DNA (grey). Scale bar = 5 μm. (**B**) Phylogenetic profiles of proteins involved in mitosis. Filled and empty circles indicate presence and absence of proteins respectively. The panels and circles are coloured to reflect the structure they are associated with: MTOC - centriole-based (centriole) and spindle pole body (SPB, yellow), nuclear envelope (NE, blue), microtubules and spindle-associated (red) and kinetochore (KT, green) (materials and methods). In addition to Ichthyosporea (*Sarc* and *Cper*) and Corallochytrea (*Clim*), profiles of key species are represented including *Homo sapiens (Hsap*), *Schizosaccharomyces pombe (Spom*), the choanoflagellate *Salpingoeca rosetta* (*Sros*), the filasterean *Capsaspora owczarzaki* (*Cowc*), the early branching chytrid fungus *Spizellomyces punctatus* (*Spun*), and the amoebozoan *Physarum polycephalum* (*Ppol*). (**C**) *C. perkinsii* has a centriolar MTOC. Single slices from TEM tomography of *C. perkinsii* cells showing top and side view through centrioles. Scale bar = 200 nm. (**D**) *S. arctica* has an acentriolar MTOC. Single slice from TEM tomography of *S. arctica* interphase nucleus. Side and top view of segmentation of *S. arctica* MTOC from an interphase nucleus. Scale bar = 500 nm.

The Opisthokonta, a major eukaryotic group that includes animals, fungi, and their deep-branching relatives, present an ideal context for such an evolutionary cell biology analysis, with species in the clade exhibiting a broad range of genome organisation modes, physiology and ecology (*21*– *23*). Importantly, either open or closed mitosis is dominant within the major animal and fungal lineages, respectively (*1, 2*). We know little about mitosis in the deep-branching opisthokonts that lie between animals and fungi, including the Choanoflagellatea, Filasterea, Ichthyosporea and Corallochytrea (Fig. 1A) (*21, 24*). Among these, Ichthyosporea, consisting of two main lineages, Dermocystida and Ichthyophonida (Fig. 1A), exhibit diverse life cycles featuring a mixture of fungal-like traits and transient multicellular stages reminiscent of early animal development (*21, 25, 26*). Most Ichthyosporea proliferate as coenocytes, multinucleated cells formed through sequential rounds of mitosis without cytokinesis, that eventually complete their life cycle through coordinated cellularization (*25*). However, a few understudied species undergo nuclear division with coupled cell cleavages (palintomic division) (*27, 28*), providing a unique opportunity to assess if and how mitotic strategies in Ichthyosporea might be linked to distinct uni- or multinucleated life cycles.

### Comparative genomics predicts divergence of the mitotic machinery within Ichthyosporea

We focussed on a set of conserved protein families (Data S1) involved in structural changes of the NE, chromosomes and spindle during mitosis (Fig. 1B and fig. S1). We identified putative orthologs and inferred gene trees of these key regulators across a set of representative opisthokonts, centred on the two major lineages of Ichthyosporea, the Ichthyophonida and Dermocystida, and using the Amoebozoa as a neighbouring outgroup that includes the model slime moulds *Dictyostelium* and *Physarum* (*29*) (Fig. 1B and fig. S1). Our results highlight features shared by all Ichthyosporea, such as the kinetochore-localised Dam complex also present in most fungi, (*30, 31*) as well as features restricted to specific ichthyosporean lineages. In an example of the latter, Ichthyophonida, including *Sphaeroforma arctica* (Fig. 1B), lack centriolar components such as Sas6 and Plk4, typical animal nucleoskeletal components such as lamins, as well as parts of the constitutive centrome associated network (CCAN) network that is otherwise broadly distributed across opisthokont genomes (Fig. 1B and fig. S1). In contrast, the dermocystid *Chromosphaera perkinsii* displays an animal-like repertoire of mitosis-related components, including a centriolar MTOC, lamins, lamin-associated proteins, and kinetochore components (Fig. 1B and fig. S1). Examining the ichthyophonid and dermocystid proteomes in greater depth predicts additional differences in spindle morphology and NPC dynamics between the two groups. For example, *S. arctica* appears to be missing several subunits of the Augmin complex responsible for nucleating new microtubules on existing spindle microtubule bundles (*32*). Although all Ichthyosporea appear to possess a PRC1/Ase1 spindle crosslinker (*33, 34*), the gene tree suggests that the *S. arctica* PRC1 is positioned on one side of an ancient duplication, clustering away from typical animal and fungal PRC1 while *C. perkinsii* retains both orthologs (fig. S2A). The conserved nucleoporin ELYS is a key regulator of post-mitotic NPC assembly, thought to be dispensable for pathways of interphase assembly and in systems with a closed mitosis. *C. perkinsii* possesses a full-length ELYS ortholog, while the truncated *S. arctica* ELYS is reminiscent of the fission yeast protein (fig. S2B and C). These analyses produce several experimentally verifiable predictions, including the presence of centrioles in Dermocystida and acentriolar MTOCs in Ichthyophonida (*25, 35, 36*), accompanied by an overall divergence in spindle architecture, NE and NPC dynamics, and kinetochore organisation between the two lineages.

### The *C. perkinsii* centriole and a novel MTOC in *S. arctica*

We first set out to examine the MTOCs of *C. perkinsii*, the only free-living dermocystid isolated to date, and *S. arctica,* the best-studied ichthyosporean model, using Transmission Electron Microscopy (TEM) and Focussed Ion Beam Scanning Electron Microscopy (FIB-SEM). As predicted by the phylogenetic analysis, we identified centrioles in *C. perkinsii* with canonical 9-fold symmetry and a diameter (220 ± 14 nm, n = 17) (Fig. 1C) very close to that of the typical animal centriole (*37*). In contrast, we find that the *S. arctica* MTOC is a multi-layered structure positioned at the outer nuclear membrane during interphase (Fig. 1D and movie S1), reminiscent of the fungal spindle pole body (SPB) (*23, 38*). During mitosis, the localisation of the MTOC shifts from the outer to the inner nuclear membrane (fig. S3A), predictive of an intranuclear spindle. A single nuclear pore was located directly underneath a subset of interphase MTOCs (fig. S3A and movie S1), possibly an intermediate in an insertion-extrusion cycle of the type best characterised in the fission yeast *Schizosaccharomyces pombe* (*19*). The presence of an animal-like centriole-based MTOC in *C. perkinsii* (Fig. 1B, C and fig. S1) and a unique NE-associated acentriolar MTOC in *S. arctica* with some fungal features (Fig. 1B, D and fig. S3), validate our combined phylogenetic and comparative cell biology approach, leading us to focus on the underlying mitotic strategies in the two species.

### Fungal-like closed mitosis in *S. arctica*

*S. arctica* proliferates through synchronised rounds of nuclear divisions without cytokinesis, resulting in the formation of multinucleated coenocytes that later undergo actomyosin-dependent cellularisation driven by the nuclear-to-cytoplasm ratio (*25, 36*). We investigated the architecture of the MTOC, microtubule cytoskeleton, NE and NPCs using TEM, FIB-SEM, and immunofluorescence (IF) in synchronised *S. arctica* cells. Classical IF protocols are ineffective in Ichthyosporea, primarily due to the presence of a thick cell wall of unknown composition (36±19% of cells stained in typical experiments, with many cells deforming following permeabilization, fig. S4A). To overcome this challenge, we implemented Ultrastructure Expansion Microscopy (U-ExM), dramatically improving both the staining efficiency (92±6% of cells stained, fig. S4A) and the spatial resolution of IF images through 4-fold isotropic expansion (fig. S4B). U-ExM revealed a prominent polar MTOC-nucleated astral microtubule (MT) network in both interphase and mitotic *S. arctica* cells (Fig. 2A). We reconstructed the spatiotemporal dynamics of *S. arctica* mitosis by ordering nuclei by spindle length along a pseudo-timeline (Fig. 2B-D, movie S2). We observe intranuclear spindle halves initially coalescing to reduce the distance between the two MTOCs (Fig. 2B-D and movie S3 and S5). During anaphase, the spindle, now linear and bundled, elongates to separate both DNA masses with minimal apparent chromosome condensation (Fig. 2B-D, movie S4). Using pan protein-labelling, MAb414 which targets NPCs and electron tomography (ET), we found that anaphase nuclei take on the characteristic dumbbell shape commonly observed in amoebozoan and fungal closed mitosis (Fig. 2D and F, movie S4 and S6), with NPCs maintaining their localisation patterns and density throughout mitosis. Remarkably, we observed a radial arrangement of the NPCs surrounding the MTOCs (fig. S5A-C). These radial arrays, also present in other *Sphaeroforma* species (fig. S5D), align along cytoplasmic MTs emerging from the MTOCs and reorganise upon MT depolymerization using carbendazim (MBC) (fig. S5E). Mild MBC treatment causes the central spindle to collapse (fig. S6A) but a small number of NE-adjacent astral microtubules persist (fig. S6A-D), indicating that the putative MT-NPC interaction at the NE might enhance microtubule stability. We confirm using TEM and FIB-SEM that NE integrity remains intact throughout mitosis (Fig. 2F, movie S5 and S6). Together, phylogenetic analysis supported by ultrastructure imaging demonstrates that the coenocytic life cycle of *S. arctica* is accompanied by closed mitosis mediated by a unique acentriolar MTOC reminiscent of fungi.

**Figure 2.**
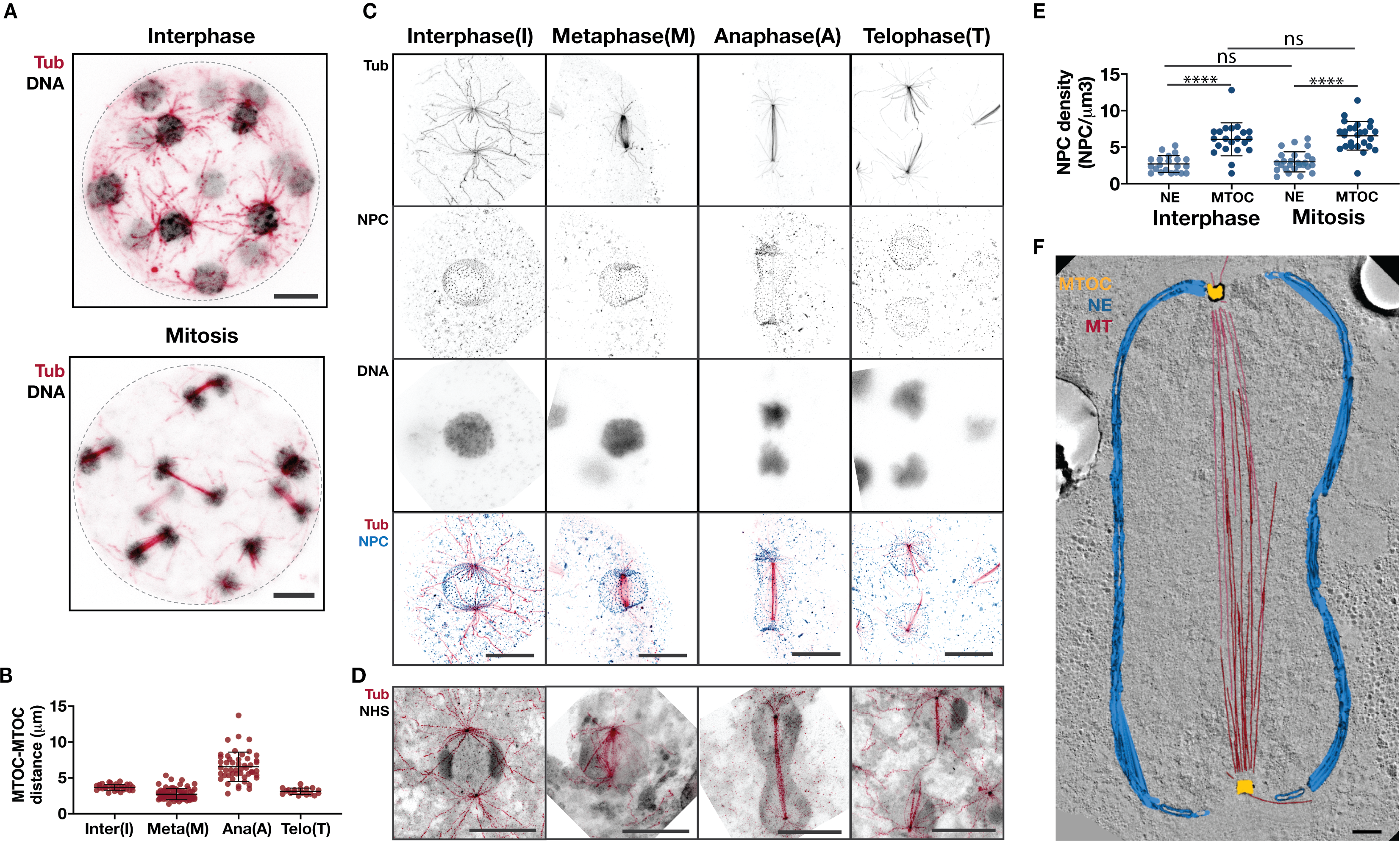
*S. arctica* undergoes closed mitosis. (**A**) Synchronised *S. arctica* coenocytes during interphase (top) and during mitosis (bottom) labelled for tubulin (red) and DNA (grey) display prominent polar astral microtubules. Scale bar = 5 μm. (**B**) Dot plot showing MTOC-MTOC distance through the different stages of the cell cycle, interphase (I) (n=29), metaphase (M) (n=75), anaphase (A) (n=49) and telophase (T) (n=19). Note that for telophase (T), the distance was measured from the MTOC to the end of the spindle MTs after splitting in two. Scale bar = 5 μm. (*P* < 0.0001). (**C**) Mitotic dynamics of microtubules (red), DNA (grey) and NPCs (blue) in *S. arctica*. Note that *S. arctica* forms intranuclear spindles during mitosis. Images shown are maximum intensity projections. Scale bar = 5 μm. (**D**) *S. arctica* carries out closed mitosis forming the characteristic dumbbell shaped nucleus in anaphase. Maximum intensity projections of nuclei with pan protein labelling (NHS ester, grey) and tubulin labelling (red). Scale bar = 5 μm. (**E**) NPC density is maintained throughout the life cycle. Dot plot showing NPC density in interphase and mitosis, measured around the MTOC and rest of the NE (n_interphase_ = 21, n_mitosis_ = 29) (ns-not significant; \*\*\*\**P* < 0.0001). (**F**) Single slice from an electron tomogram of dumbbell-shaped *S. arctica* anaphase nucleus overlaid with a 3D model showing the intranuclear spindle (red), MTOC (yellow) and NE (light blue). Scale bar = 500nm

### Animal-like open mitosis in *C. perkinsii*

We then investigated mitosis in the dermocystid *C. perkinsii*, which we initially predicted to possess an animal-like complement of mitotic regulators (Fig. 1B) and which, like its close parasitic relative *Sphaerothecum destruens* (*27*) proliferates through palintomic divisions or cell cleavages (Fig. 3A). Reconstructing the sequence of *C. perkinsii* mitotic stages reveals a strikingly human-like spindle nucleated from centrioles (Fig. 3B, movie S7 and S8), with several key structural similarities including the presence of kinetochore fibres and bridging fibres, scaling of inter-kinetochore distances and twist of the spindle (Fig. 3C-F, fig. S7 and Supplementary text), an equatorial arrangement of condensed chromosomes in metaphase (Fig. 3B), and the simultaneous segregation of chromosome complements to opposing poles in anaphase (Fig. 3B). In contrast to *S. arctica* nuclear division, *C. perkinsii* mitosis is characterised by NE breakdown and reassembly kinetics typical of mitosis in human cells (Fig. 3B, F and movie S7-9), with NPCs disappearing in prophase accompanied by a loss of NE integrity (Fig. 3B and 3C) and reappearing in telophase (Fig. 3B). Given the accumulating evidence suggesting a sharp divergence in mitosis between dermocystids and ichthyophonids, we examined a range of additional ichthyosporean species and their closest living relative, the corallochytrean *Corallochytrium limacisporum* (*39*). Using U-ExM of MTs, NE and NPCs, we find that other coenocytic ichthyophonids (three distinct *Sphaeroforma* species as well as *Creolimax fragrantissima*), exhibit a spindle architecture reminiscent of *S. arctica* and appear to undergo a comparable closed mitosis (fig. S8). In contrast and in line with our initial predictions based on the phylogenetic analysis (Fig. 1A and fig. S1), *C. limacisporum*’s uninucleate life cycle (*40*) appears to be facilitated by open mitosis (fig. S8). The presence of an animal-like open mitosis in early-diverging holozoan species (*C. perkinsii and C. limacisporum*) suggests that this mitotic strategy predates the emergence of animals.

**Fig. 3.**
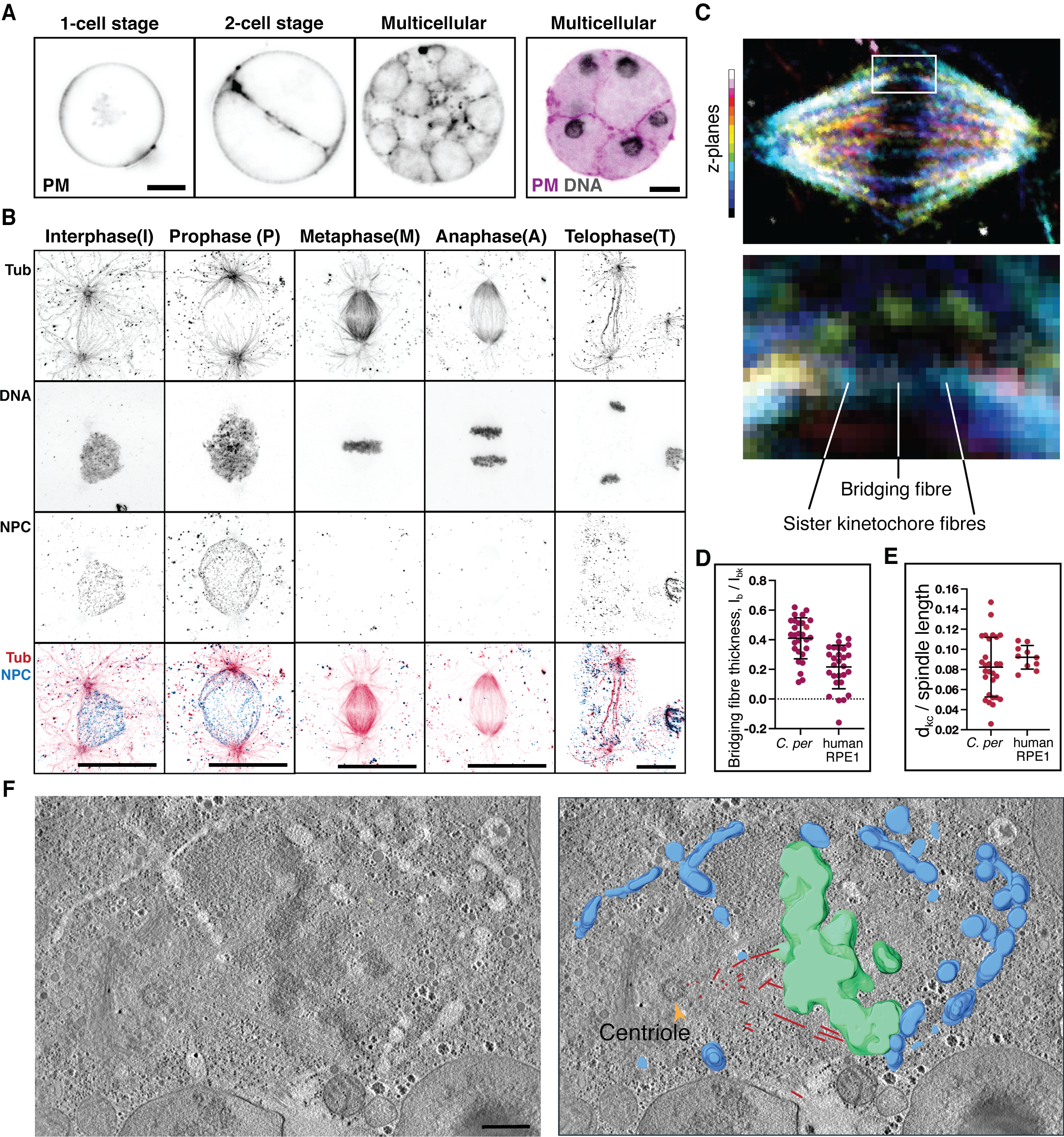
*C. perkinsii* undergoes open mitosis. (**A**) Cell cleavages shown in *C. perkinsii* cells at different life cycle stages labelled for plasma membrane (PM, purple) and DNA (grey). Scale bar = 5 μm. (**B**) Maximum intensity projections of *C. perkinsii* nuclei at different stages of the cell cycle, namely interphase(I), prophase(P), metaphase(M), anaphase(A) and telophase(T), in cells labelled for tubulin (red), DNA (grey) and NPCs (blue). Scale bar = 5 μm. (**C**) *C. perkinsii* spindle in metaphase coloured for depth as shown in the legend on the left (top); enlarged section of the spindle showing kinetochore and bridging fibres (bottom). (**D**) Bridging fibre thickness relative to the kinetochore fibre thickness. (**E**) Interkinetochore distance scaled to the spindle long axis. Data for RPE1 cells in (**D**) and (**E**) was taken from Štimac et al., 2022 (*32*) (**F**) Single slice from an electron tomogram of *C. perkinsii* mitotic nucleus overlaid with a 3D model showing the spindle MTs (red), chromosomes (green) and NE (blue). The yellow arrowhead indicates the centriole in focus. Scale bar = 500 nm.

## Discussion

Despite the fundamental role of cell division in propagating cellular life, the diversity of eukaryotic mitotic mechanisms, and in particular the mode of NE remodelling and design principles of the spindle, have been challenging to study. This is mainly due to the fact that the only open and closed mitotic processes that are understood in significant molecular detail derive from animals and fungi. Here we provide evidence for a switch between open and closed mitosis within Ichthyosporea (Fig. 4), cementing their role as a key group of species, along with other deep-branching Holozoa, for investigating the evolution of mitosis. The striking ultrastructural similarities between *C. perkinsii* and animal mitosis on one hand (Fig. 3) and *S. arctica* and fungal mitosis on the other (Fig. 2), strongly support the idea of a tight coupling between NE remodelling and architectural features of the spindle, chromosomes and NE. Genomic and imaging analysis of the closest known ichthyosporean relatives, the Corallochytrea, suggests that the ancestral ichthyosporean possessed a centriole and was capable of open mitosis. These features are likely retained in the dermocystids to support a flagellated life cycle stage (*28, 39*) and, as we show in this study for *C. perkinsii*, palintomic divisions through open mitosis (Fig. 3); features that were therefore likely inherited by the ancestor of animals. This implies in turn that the adoption of an exclusively coenocytic life cycle within the Ichthyosporea was accompanied by the loss of the centriole, the *de novo* evolution or retention of a distinct, nucleus-associated MTOC, and a switch to closed mitosis. The broad range of examples of closed mitosis with centrioles (Fig. 4) (*41–43*) and open mitosis without centrioles (*44, 45*) argues that the presence of a flagellum does not constrain the mode of mitosis. Instead, our results suggest that having a coenocytic life cycle stage, in which more than two nuclei must divide and faithfully segregate in a shared cytoplasm, requires a closed or semi-closed mitosis. This model is supported by data on the semi-closed mitosis of the *Drosophila* coenocytic embryo (*46, 47*), the germline of various animal lineages (*48–50*), the closed or semi-closed mitosis associated with hyphal growth in fungi (*7, 51*), and the switch between open and closed mitosis in the complex life cycle of the amoebozoan *Physarum polycephalum* (*52, 53*). A corollary of this hypothesis is that once closed mitosis has evolved, it can persist even if the organism returns to a unicellular, uninucleate life cycle (*6, 54*) but in such cases is apparently no longer under strict selection to remain closed (*55, 56*).

**Fig. 4.**
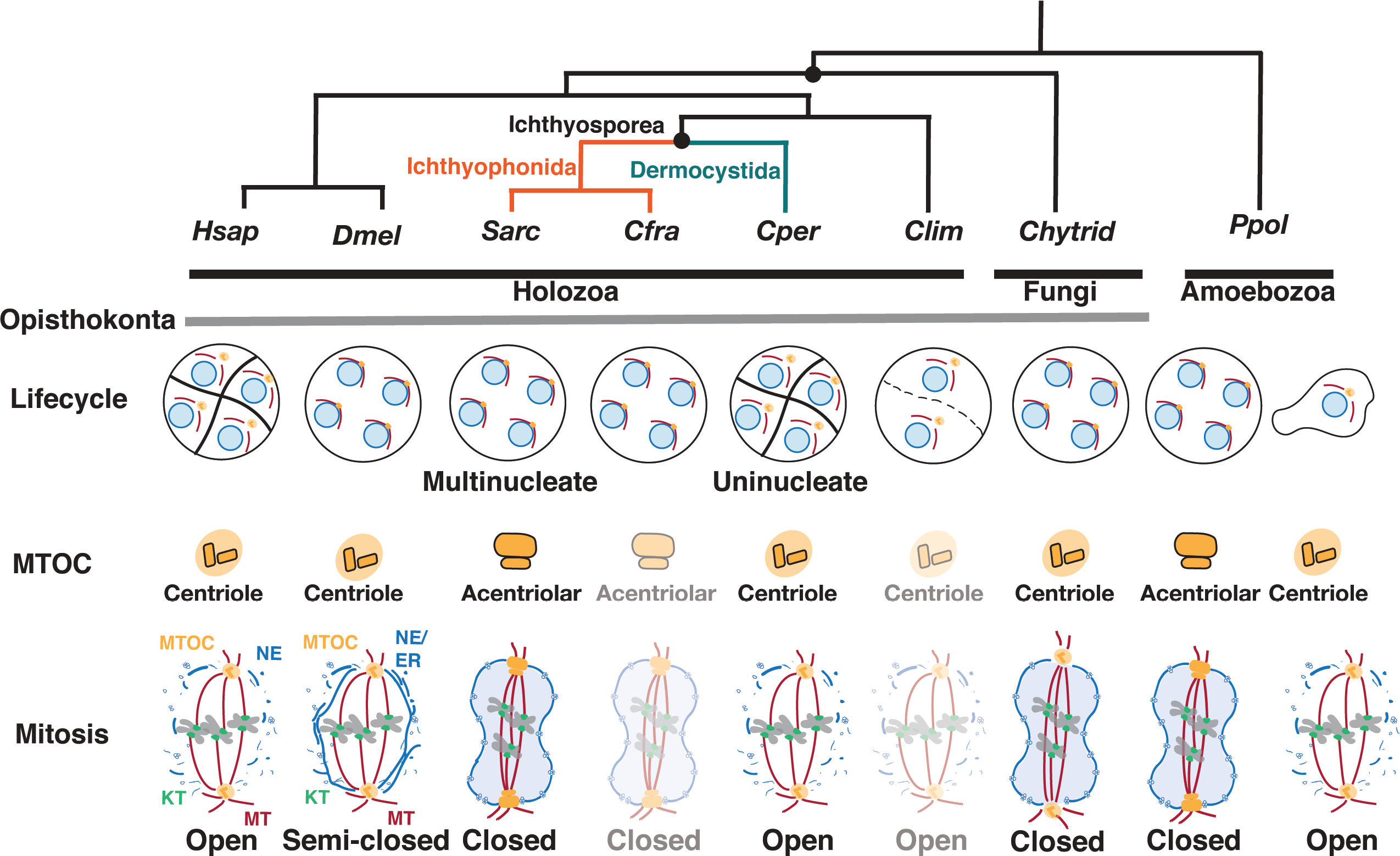
Specialisation of mitotic strategies coupled to distinct life cycles in opisthokonts. Ichthyophonida, including the model ichthyosporean, *S. arctica* (*Sarc*) and *C. fragrantissima* (*Cfra*) form multinucleated cells, like many fungal and amoebozoan species (*Ppol*), through a series of closed mitoses. As in classical fungal closed mitosis, ichthyosporean closed mitosis involves NE embedding of the acentriolar MTOC and remodelling of the nucleus into dumbbells upon elongation of the intranuclear spindle Dermocystids, on the contrary, divide by cell cleavages resulting in uninucleated cells and undergo centriole-mediated open mitosis with NE breakdown involving spindle architecture reminiscent of mammalian (*Hsap*) mitosis. The corallochytrean *C. limacisporum*, which divides as uninucleated cells through open mitosis, suggests that open mitosis is ancestral in Holozoa. In coenocytic insect embryos *(DmeI)*, the integrity of the nuclear compartment is maintained by the ER. Across opisthokonts and amoebozoan (*Ppol*) outgroups, the data suggests a coupling between open or closed mitosis with uni- and multinucleated life cycles.

Beyond mitosis, our work here provides a generalisable framework for evolutionary cell biology, linking comparative genomics to cell biology across a range of non-model species. A key challenge in mapping genotype to phenotype is the mismatch of scales: we have access to many more high-quality genomes than experimental model systems, and developing a new species into a model system demands many years of dedicated effort. Here we bridge that gap using a volumetric ultrastructure imaging approach that combines the scalable tools of expansion microscopy with the traditional advantages of EM. When guided by phylogenetics, such a framework can enable a comparative cell biology approach to investigating major evolutionary transitions.

## Acknowledgments

Sophie Martin, Pierre Gönczy, Jan Ellenberg, Buzz Baum, Rebecca Heald, Agathe Chaigne, Pedro Pereira, Siân Culley, Ishier Raote, Thomas Quail, Hanh Vu, Flora Vincent, Anna Erzberger, Jenna Elliot, Nicoletta Petridou, Niccolo Banterle, Michael Dorrity and all members of the Dey, Dudin and Schwab labs for comments on manuscript and general feedback; Hiroshi Suga for *C. limacisporum* and *C. perkinsii* cultures. We thank the Advanced Light Microscopy Facility (ALMF) at the European Molecular Biology Laboratory (EMBL), Carl Zeiss, the Electron Microscopy Core Facility (EMCF) at EMBL and the biological electron microscopy facility at EPFL for their support.

## Funding

OD and MO are funded by an Ambizione fellowship from the Swiss National Science Foundation (PZ00P3_185859). GD, HS, CB, PR and YS are funded by the European Molecular Biology Laboratory. H.S. is supported by the EMBL Interdisciplinary Postdoctoral Fellowship (EIPOD4) programme under Marie Sklodowska-Curie Actions Cofund (grant agreement number 847543). E.T. is supported by an NWO-Veni Fellowship (VI.Veni.202.223). M.T. has been supported by the ‘‘Young Researchers’ Career Development Project—training of doctoral students’’ of the Croatian Science Foundation. I.M.T. acknowledge the support of European Research Council (ERC Synergy Grant, GA 855158), Croatian Science Foundation (HRZZ project IP2019-04-5967), and projects co-financed by the Croatian Government and European Union through the European Regional Development Fund—the Competitiveness and Cohesion Operational Programme: QuantiXLie Center of Excellence (grant KK.01.1.1.01.0004) and IPSted (grant KK.01.1.1.04.0057).

## Author contributions

Conceptualization: HS, OD, GD

Formal Analysis: HS, MO, CB, MT, ET, PR, YS, OD, GD

Funding Acquisition: IMT, YS, OD, GD Methodology: HS, MT, ET, IMT, PR, YS, OD, GD

Project administration and supervision: YS, OD, GD Visualization: HS, OD, GD

Writing - Original draft: HS, OD, GD

Writing - review and editing: HS, MO, CB, MT, ET, IMT, PR, YS, OD, GD

## Competing interests

Authors declare that they have no competing interests.

## Data and materials availability

Data associated with the figures is available at https://doi.org/10.6084/m9.figshare.c.6639812.v1.

## Supplementary Materials

Materials and Methods

Supplementary text

Tables S1

Movies S1 to S9

Data S1

## Supplementary Materials

### Materials and Methods

#### Phylogenetic analysis

We generated profiles of representative proteins from mitosis-associated cellular components including the centrosome, spindle pole body, spindle, nuclear envelope and the kinetochore for early branching animal and fungal lineages alongside model species with well characterised mitosis. The proteomes of *Homo sapiens, Salpingoeca rosetta, Monosiga brevicollis, Capsaspora owczarzaki, Schizosaccharomyces pombe, Spizellomyces punctatus, Dictyostelium discoideum* and *Physarum polycephalum* were obtained from Eukprot v02.2020_06_30 (*57*). The ichthyosporean proteomes were obtained from recent phylogenomic studies (*23, 25, 39*). To identify putative orthologs we first searched for homologs with human proteins using phmmer (HMMER 3.3.2, Nov 2020; http://hmmer.org/) (*58*). In case of divergence or absence of human proteins, searches were also carried out with proteins from other model species including *S. pombe, D. melanogaster* and *D. discoideum*. This was followed by Hidden Markov model (HMM)-based searches with the associated PFAM (http://pfam.wustl.edu) (*59*) models using hmmsearch (HMMER 3.3.2, Nov 2020; http://hmmer.org/) (*58*). For proteins where homologs were not recovered by existing HMMs, new HMMs were generated. The multiple sequence alignment was done with MAFFT v7.490 using linsi optimised for local homology (*60*). The alignments were inspected and trimmed using TrimAl v1.2 to remove the less conserved regions (*61*). The trimmed alignments were used for tree inference with IQTree v2.0.3 2020 using the model finder and ultrafast bootstraps (1000) bootstraps (*62–64*). The trees were visualised and annotated using FigTree (Molecular evolution, phylogenetics and epidemiology, Andrew Rambaut, http://tree.bio.ed.ac.uk/software/figtree/). This process was performed iteratively to obtain better alignments that gave trees with higher bootstrap values. The alignment was then used to generate an HMM using the hmmbuild command in HMMER. This process was performed iteratively while incorporating the newly discovered homologs in the next round. The protein sequences are provided as fasta files. For most kinetochore proteins, iterative similarity searches were performed using previously generated HMMs of which candidate genes were scrutinised based on known domain and motif topologies (*17*).

#### Culture conditions

*S. arctica* cultures were maintained at 17°C in Marine broth (MB, Difco, 37.4g/L) and synchronised as previously described (*25, 35, 36*). Briefly, for synchronisation, 1/16 MB was prepared by diluting MB in artificial seawater (Instant Ocean, 37g/L). Cultures were diluted 1:100 in 1/16 MB and grown for 3 days to obtain saturated cultures. To obtain a synchronised culture, the saturated cultures were inoculated 1:50 in fresh MB. To obtain the 8-32 nuclear stage, cells were fixed around 28.5 hours post inoculation. Other *Sphaeroforma sp., S. gastrica, S. nootkatensis* and *S. napiecek,* and *Creolimax fragrantissima* were maintained at 17°C in MB similar to *S. arctica*. *C. perkinsii* and *Corallochytrium limacisporum* were grown at 23°C protected from light. For maintenance, cultures were diluted 1:1000 every two weeks and restarted from a cryopreserved stock every 6 months. For experiments, 6 day old cultures were filtered using a 5µm filter to obtain small newborn cells which are then diluted 1:100 in *C. perkinsii* medium to obtain synchronous cultures. The cells were fixed at the 1-8 cell stage (60-90 hrs post dilution) in order to capture the initial mitotic events.

#### Immunostaining

The cell culture flasks were scraped and the suspension was added to 15ml falcons to sediment for 15-30 mins. The supernatant was removed and cells were transferred to 1.5ml microfuge tubes and fixative was added for 30 mins. The cells were fixed with 4% formaldehyde (FA) in 250mM Sorbitol solution, washed twice with 1X phosphate buffer saline (PBS) and resuspended in 20-30μl PBS. Cells were permeabilized using nine freeze-thaw cycles (liquid N_2_,10 s: 42°C, 1 min). This was followed by blocking in 3%BSA in PBST (1X PBS with 0.1% Tween20). Primary antibody (Tubulin-E7 antibody DSHB, NB600-936 Novus Biologicals, AA344 and AA345 (ABCD antibodies and anti-nuclear pore complex proteins - MAb414 Biolegend 902901) was used at 1:500 to 1:1000 and incubated at 4°C overnight or 2-5 hrs at 37°C. This was followed by 3 washes for 10 mins at RT and addition of the secondary antibody. Incubation was done at 4°C overnight or 2-5 hrs at 37°C. The cells were then washed and resuspended in fresh 1X PBS for imaging. DNA was stained with Hoechst 33352 at a final concentration of 0.4 µM. For live cell imaging cells were stained with FM-464 at a final concentration 10 μM.

#### Ultrastructural Expansion Microscopy (U-ExM)

U-ExM was performed as previously described (*65*). Briefly, the cells were fixed with 4% FA in 250mM Sorbitol solution, washed twice with 1X PBS and resuspended in 20-30μl PBS. The fixed cells were then allowed to attach to 12 mm poly-l-lysine coated coverslips for 1 hr. This was followed by anchoring in AA/FA (1% Acrylamide (AA)/ 0.7% Formaldehyde (FA)) solution for at least 5 hrs and up to 12 hrs at 37°C. A monomer solution (19% (wt/wt) sodium acrylate (Chem Cruz, AKSci 7446-81-3), 10% (wt/wt) Acrylamide (Sigma-Aldrich A4058), 0.1% (wt/wt) N, N’-methylenbisacrylamide (Sigma-Aldrich M1533) in PBS) was used for gelation and gels were allowed to polymerize for 1 hr at 37°C in a moist chamber. For denaturation, gels were transferred to the denaturation buffer (50 mM Tris pH 9.0, 200 mM NaCl, 200 mM SDS, pH to 9.0) for 15 min at RT and then shifted to 95°C for 1 hr. Following denaturation, expansion was performed with multiple water exchanges as previously described (*65*). Post expansion, gel diameter was measured and used to determine expansion factor. For all U-ExM images, scale bars indicate actual size; rescaled for gel expansion factor. Pan-labelling of U-ExM was done at 1:500 with Dylight 405 (Thermo Fischer, 46400) or Alexa Fluor NHS-Ester 594 (Thermo Fischer A20004) in 1x PBS or NaHCO_3_ for 1.5 hrs. Immunostaining was performed as mentioned above. All antibodies were prepared in 3% PBS with 0.1% Tween 20.

#### Light Microscopy

For immunolabelled cells, we used Poly-l-Lysine coated Ibidi chamber slides (8 well - Ibidi 80826). The wells were filled with 1X PBS and 0.4 µM Hoechst 33342 (Thermo Fischer 62249) was added. Immunostained cells were added to wells and allowed to settle for an hour before imaging. Imaging was done on the Zeiss LSM 880 using the Airyscan Fast mode using the Plan-Apochromat 63x/1.4 Oil DIC M27 Objective. For staining efficiency, sample overviews were imaged in LSM mode using the tilescan function with the Plan-Apochromat 63x/1.4 Oil DIC M27 objective. For immunolabeled U-ExM gels, we also used Poly-l-Lysine coated Ibidi chamber slides (2 well - Ibidi 80286, 4 well - Ibidi). Gels were cut to appropriate size to fit the Ibidi chambers and added onto the wells. The gels were overlaid with water to prevent drying or shrinkage during imaging. The gels were either imaged using the Zeiss LSM 880 with the Airy fast mode using a Plan-Apochromat 63x/1.4 Oil DIC M27 objective or using an upright Leica SP8 confocal microscope with a HC PL APO 40X/1.25 Glycerol objective.

#### Image analysis

For Immunostaining efficiency measurements (fig. S4A), immunostained (IF) and expanded (U-ExM) samples were stained for microtubules (E7 antibody, DSHB). Cells were imaged using confocal microscopy in tilescan mode. Hoechst 33342 or NHS-ester was used as a reference to determine the percentage of immunostained cells.

For NPC density measurements (Fig. 2E), NPC density was determined using nuclei from MAb414 (Biolegend 902901) labelled U-ExM gels. In *S. arctica* NPC densities are different around the MTOC as compared to the rest of the nucleus. Thus, two ROIs were selected per nucleus, one within the radial arrays in the vicinity of the MTOC and a second one away from it (marked in the graph as NE). Each ROI was a 5µm cube. The nuclei were classified as interphase or mitotic, based on nuclear shape and presence or absence of intranuclear MTs. The images were thresholded and binarized. The 3D Object Counter plugin was used to obtain NPC counts. The counts were divided by cube volume to obtain NPC density. The measurements were corrected for the gel expansion factor to obtain actual NPC density per µm^3^.

For spindle pole body dimensions (fig. S3B), analysis of SPB dimension was done using pan-labelled U-ExM gels. The images were cropped to a 5μm region around the SPB, thresholded and binarized. The ‘Analyse particle’ function was used to obtain SPB shape measurements including width and height (fig. S3B). The measurements were corrected for gel expansion factor to obtain actual width and height.

For *S. arctica* MTOC-MTOC distance (Fig. 2B), the distance was determined using Tubulin labelled U-ExM gels. The images were thresholded using morphological filtering and binarized. The structure was then skeletonized and the Analyse Skeleton plugin (https://github.com/fiji/AnalyzeSkeleton) was used to determine spindle length. In cases where this was not possible, MTOC positions were marked manually and Euclidean distance was calculated between the two points.

*C. perkinsii* centriole diameter was measured in Fiji from TEM tomography images. Both longitudinal and transversely placed centrioles were used for the analysis. Centrioles were placed at varying angles to the sectioning and imaging plane.

Image analysis was performed using Fiji software (*66, 67*). All figures were assembled with Illustrator 2022 CC 2020 (Adobe). Graphs were generated using GraphPad Prism 9. 3D reconstructions of movies S3, 4, 7 and 8 were done in Imaris v992.

#### Live cell imaging

Light-sheet microscopy in movie S2 was performed using the LS1 Live 246 light sheet microscope system (Viventis®) as previously described using a 25X 247 1.1 NA objective (CFI75 Apo 25XW; Nikon) and an sCMOS camera (Zyla 4.1, Andor) (*36*). Light-sheet imaging was conducted in a room specifically cooled at 17°C using an air-conditioning unit.

#### Electron Microscopy

##### Sample preparation 1

For TEM tomography of *S. arctica* and *C. perkinsii* cells, samples were concentrated by sedimentation and high pressure frozen with the HPM010 (Abra Fluid) using 200 μm deep, 3 mm wide aluminium planchettes (Wohlwend GmbH). Freeze substitution was done using the AFS2 machine (Leica microsystems) in a cocktail containing 1% OsO_4_, 0.1% uranyl acetate (UA) and 5% water in acetone. The samples were incubated as follows: 73h at −90°C, temperature increased to −30°C at a rate of 5°C/h, 5h at −30°C, temperature increased to 0°C at a rate of 5°C/h, 4 x 0.5h rinses in water free acetone at 0°C. This was followed by Epon 812 (Serva) infiltration without BDMA (25% – 3h at 0°C, 50% overnight at 0°C, 50% - 4 h at room temperature (RT), 75% - 4h, 75% overnight, 100% 4h (x2) and 100% overnight. This was followed by exchange with 100% Epon 812 with BDMA (4h x 2, followed by overnight). After this the samples polymerised in the oven at 66°C for over 2 days. The samples were then cut using an ultramicrotome (Leica UC7) in 70nm sections screened by 2D TEM (Jeol 1400 Flash) to assess sample preparation. For TEM tomography (300 nm sections), sections were post-stained with 2% UA in 70% methanol (5 min, RT) and in Reynolds lead citrate (2 min, RT).

##### Sample Preparation 2

For serial tomography and FIB-SEM of *S. arctica* cells, samples were concentrated and high pressure frozen as mentioned above. Freeze substitution was done in the AFS2 machine (Leica microsystems) in a cocktail containing 1% OsO_4_, 0.5% uranyl acetate (UA) and 5% water in acetone. The samples were incubated as follows: 79h at −90°C, temperature increased to −60°C at a rate of 2°C/h, 10h at −60°C, temperature increased to −30°C at a rate of 2°C/h,10h at −30°C, temperature increased to 0°C at a rate of 5°C/h, 1h at 0 °C. After this, the samples were rinsed in acetone and further incubated in 0.1% thiocarbohydrazide 10% water in acetone for 30min at room temperature, followed by a 1% OsO_4_ in acetone. After rinsing, the samples were infiltrated in Durcupan ATM (Sigma) and finally polymerized in a 60°C oven for 72h. The OsO_4_ step and the infiltration were performed in a Biowave (Ted Pella, inc). The samples were then sectioned using an ultramicrotome (Leica UC7) for serial section TEM tomography (300 nm sections). The sections were post-stained with 2% UA in 70% methanol (5 min, RT) and in Reynolds lead citrate (2min, RT). Tomograms were acquired with a Tecnai F30 (Thermo Fisher Scientific) using SerialEM^39^ and reconstructed and joined with Imod Etomo. After this a 70nm section was collected and screened by 2D TEM (Jeol 1400 Flash) to target interphase and mitotic cells for FIB-SEM analysis. The samples were then mounted on a SEM stub using silver conductive epoxy resin (Ted Pella, inc), gold sputter coated (Quorum Q150RS) and imaged by FIB-SEM. The acquisition was performed using a Crossbeam 540 or 550 (Zeiss) following the Atlas 3D workflow. SEM imaging was done with an acceleration voltage of 1.5 kV and a current of 700pA using an ESB detector (1100V grid). Images were acquired at 5×5 nm pixel size and 8nm slices were removed at each imaging cycle. FIB milling was performed at 700pA current. For segmentation and visualisation we used 3DMod and Amira (version 2019.3 or 2020.1; Thermo Fisher Scientific).

#### Twist analysis

To calculate spindle twist for *C. perkinsii*, Fiji Software (ImageJ, National Institutes of Health, Bethesda, MD, USA) (*66*) was used to analyse microscopy images of horizontal spindles. Only images with both spindle poles in the same plane or in two consecutive planes apart in each direction of the z-stack were included in the analysis to prevent spindle tilt from affecting the spindle twist calculation. Horizontal spindles were transformed into a vertical orientation (end-on view) using a code written in R programming language in RStudio (*68*). In the transformed stack, microtubule bundles and poles appear as blobs. The spindle poles are tracked manually using the Multipoint tool in ImageJ. Next, we used the optical flow method to calculate the twist (*69*) and presented the absolute values in the graph. The tracing of bundles and twist calculations are written in Python programming language using PyCharm IDE, with external libraries such as NumPy, scikit-image, Matplotlib, PIL, OpenCV, and SciPy. The code and instructions are available at GitLab: https://gitlab.com/IBarisic/detecting-microtubules-helicity-in-microscopic-3d-images (*67*). Twist values for RPE1 cells expressing CENP-A-GFP and centrin1-GFP were taken from Trupinić et al., 2022 (*69*).

#### Analysis of spindle length, width and interkinetochore distance

To measure spindle length, width and interkinetochore distance, the Line tool in Fiji Software (ImageJ, National Institutes of Health, Bethesda, MD, USA) (*66*) was used. Length was measured by drawing a line from pole to pole of the spindle. In *C. perkinsii* the pole positions were determined visually as the outermost points of the spindle along the central spindle axis. In RPE1 cells expressing CENP-A-GFP and centrin1-GFP, length was measured by using the images from Štimac et al., 2022 (*32*). and line was drawn from one centrosome to the other. Width in *C. perkinsii* was measured by drawing a line across the equatorial plane of the spindle, with the line ending at the outer edges of a spindle. Width in RPE1 cells expressing CENP-A-GFP and centrin1-GFP was measured by drawing a line across the equatorial plane of the spindle, with the line ending at the outer kinetochore pairs. Interkinetochore distance in *C. perkinsii* was measured by using Rectangle tool in Fiji Software which was drawn between endings of k-fibres at the spindle midzone. Interkinetochore distance in RPE1 cells expressing CENP-A-GFP and centrin1-GFP was taken from Štimac et al., 2022 (*32*). It was not possible to measure the interkinetochore distance in *S. arctica* because of the spindles’ tight microtubule bundles and it was not possible to distinguish kinetochore microtubules.

#### Analysis of the bridging fibre intensity

To measure the intensity of bridging fibres (*70*) we used Square tool (ImageJ, National Institutes of Health, Bethesda, MD, USA) (*66*). In *C. perkinsii*, the position of the square when measuring bridging fibre intensity was on the fibre located between the endings of kinetochore fibres. As we did not have labelled kinetochores and could not determine where a single kinetochore fibre is, we put squares close to the end of kinetochore fibres to obtain values of kinetochore fibres together with bridging fibres (I_bk_) (fig. S7A). Background was measured and subtracted as follows: I_b_ = I_b+bcg_ – I_bcg_ for bridging fibres and I_bk_ = I_bk+bcg_ – I_bcg_ for bridging fibres together with kinetochore fibres. The values for RPE1 cells expressing CENP-A-GFP and centrin1-GFP were taken from Štimac et al, 2022 (*69*).

#### Statistical Analysis

Results are reported as mean ± standard deviation. Statistical parameters including, the numbers of cells or nuclei analysed, n, and statistical significance are reported in the figure legends. Statistical significance was calculated by Mann-Whitney U, Kruskal-Wallis or Student *t* tests. Asterisks in graphs indicate the statistical significance (\**P* < 0.05; \*\**P*< 0.01; \*\*\**P* < 0.001; \*\*\*\**P* < 0.0001). Statistical analysis was performed in GraphPad Prism 9.

### Supplementary text

The overall architecture of MTs in *C. perkinsii’*s spindle is strikingly similar to that of human somatic cells (Fig. 3C-F and S7). However, *C. perkinsii*’s spindles are 3.5-folds shorter and 4.3-folds narrower than human retinal pigment epithelial-1 (RPE-1) (fig. S7), indicating a leaner structure. Additionally, we detect bundles of microtubules that end sharply near the equatorial plane in *C. perkinsii* spindles bearing analogy with kinetochore fibres (Fig. 3C-E). The interkinetochore distance was 4-folds smaller than in RPE-1 cells, suggesting a conserved ratio between spindle length and interkinetochore distance. Notably, near sister kinetochore fibres, we find a microtubule bundle resembling the bridging fibre in human cells, which consists of antiparallel microtubules laterally linking sister kinetochore fibres (*70*). The tubulin signal intensity of the bridging fibre, *I*_b_, was 41% of the tubulin signal intensity next to the kinetochore, represented by *I*_bk_ = *I*_b_ + *I*_k_ (fig. 3E). Using this fraction, we estimated that the number of microtubules in the bridging fibre (*I*_b_/*I*_k_ = 1/(*I*_bk_/*I*_b_-1)) is 70% of the number of microtubules in the kinetochore fibre which lies between the known range of 30% for RPE-1 cells (*32*) and 82% for the HeLa cell line^40^. In agreement with the finding that *C. perkinsii* possesses the majority of metazoan kinetochore components, we found that its spindles contain well-defined microtubule bundles that resemble kinetochore fibres, which bind to and pull-on kinetochores in spindles across various organisms ranging from yeast to humans. The distance between sister kinetochore fibres in *C. perkinsii* and human spindles was 8-9% of the spindle length. This scaling of the interkinetochore distance suggests that the molecular and biophysical mechanisms governing spindle organisation and forces may be generally conserved between these two species. In human cells, bridging fibres balance the interkinetochore tension during metaphase (*70–73*), help in preventing and correcting incorrect attachments of kinetochores to microtubules (*32, 74*), promote chromosome alignment at the spindle equator (*75, 76*), and facilitate chromosome segregation during anaphase (*77, 78*). Exploring the function of bridging fibres in crucial aspects of mitosis in *C. perkinsii* will be an intriguing area for further investigation.

**Fig. S1.**
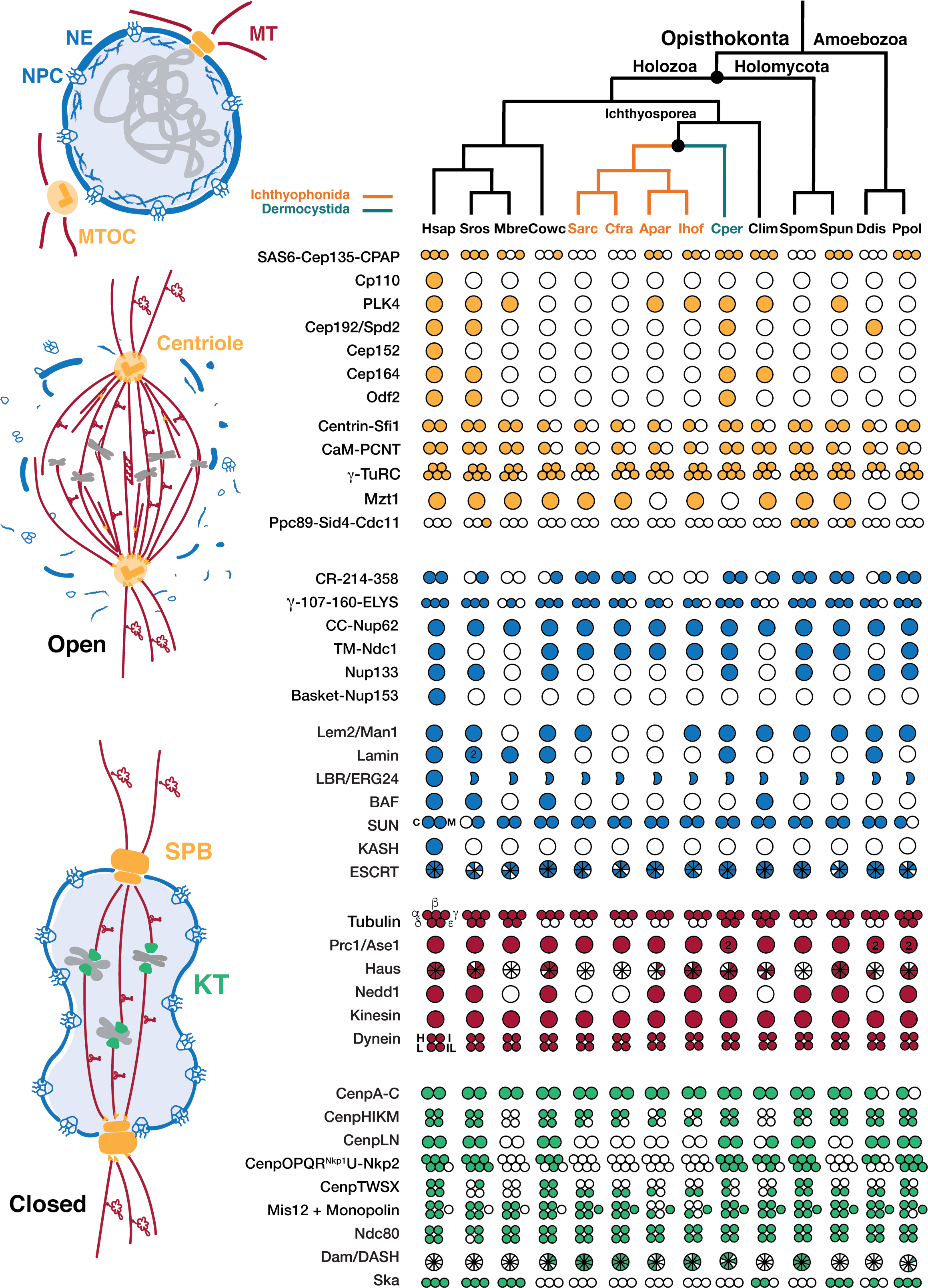
Divergence in mitosis-associated protein profile in close relatives of animals and fungi. Phylogenetic profiles of proteins involved in mitosis. Filled and empty circles indicate presence and absence of proteins, respectively. The panels and circles are coloured to reflect the structure (left) each is associated with: microtubule organising centres (MTOC (both(*4–7*) Centriole and SPB), yellow), nuclear envelope (NE, blue), microtubules and spindle-associated (MT, red), kinetochore (KT, green) and DNA/chromosomes (grey) (materials and methods). In addition to Ichthyophonida (*Sarc-S. arctica*, *Cfra-Creolimax fragrantissima, Apar-Amoebidium parasiticum* and *Ihof-Ichthyophonus hofleri)* and Dermocystida *Chromosphaera perkinsii* (*Cper*) and Corallochytrea *Corallochytrea limacisporum* (*Clim*), profiles of key species are represented including *Homo sapiens (Hsap*), *Schizosaccharomyces pombe (Spom*), the choanoflagellate *Salpingoeca rosetta* and *Monosiga brevicollis* (*Sros* and *Mbre*), the filasterean *Capsaspora owczarzaki* (*Cowc*), the early branching chytrid fungus *Spizellomyces punctatus* (*Spun*), and two amoebozoan species *Dictyostelium discoideum* (*Ddis*) and *Physarum polycephalum* (*Ppol*).

**Fig. S2.**
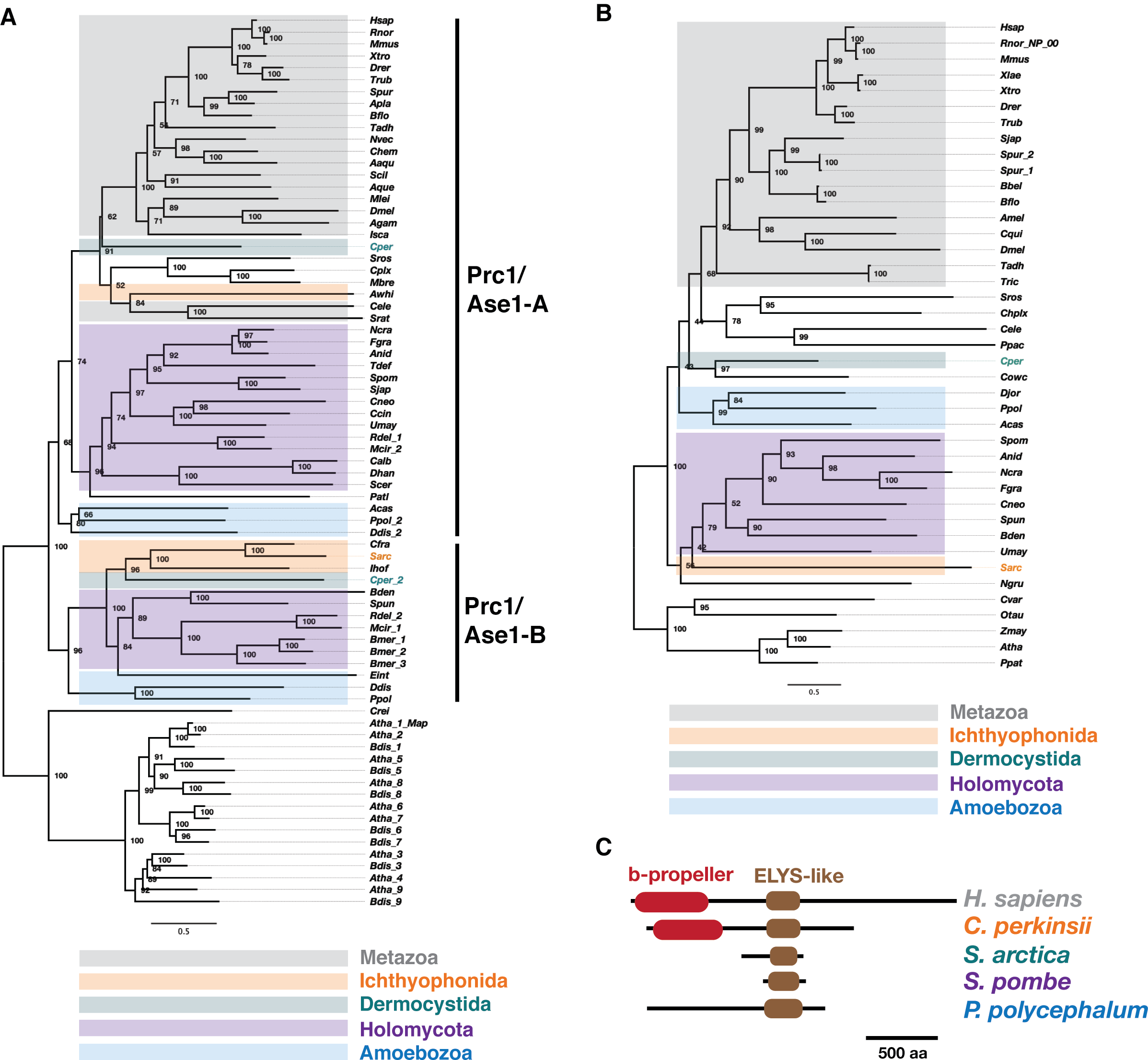
Divergence of selected mitotic proteins within Ichthyosporea. (A) In contrast to the A-type MT crosslinker protein PRC1/Ase1 present in most animal lineages, the ichthyophonid *S. arctica* retains the B-type protein, putting it on the opposite side of an early duplication. Both orthologs resulting from this duplication are found in amoebozoans, early branching fungi and the dermocystid *C. perkinsii*. Trees were made using IQtree v2.0.3 with ultrafast bootstrap (1000) using the LG+IG+G4 model. **(B)** Phylogenetic distribution of ELYS, involved in post-mitotic NPC reassembly. Trees were made using IQtree v2.0.3 with ultrafast bootstrap (1000) and LG+F+I+G4 model. The trees were visualised and annotated using FigTree (materials and methods) Numbers on the nodes are bootstrap values. Branch length units are arbitrary. **(C)** Dermocystid *C. perkinsii* has animal-like ELYS architecture with a central ELYS domain and an N-terminal ꞵ-propeller domain that has been lost in the fission yeast *S. pombe* as well as the ichthyophonid *S. arctica*.

**Fig. S3.**
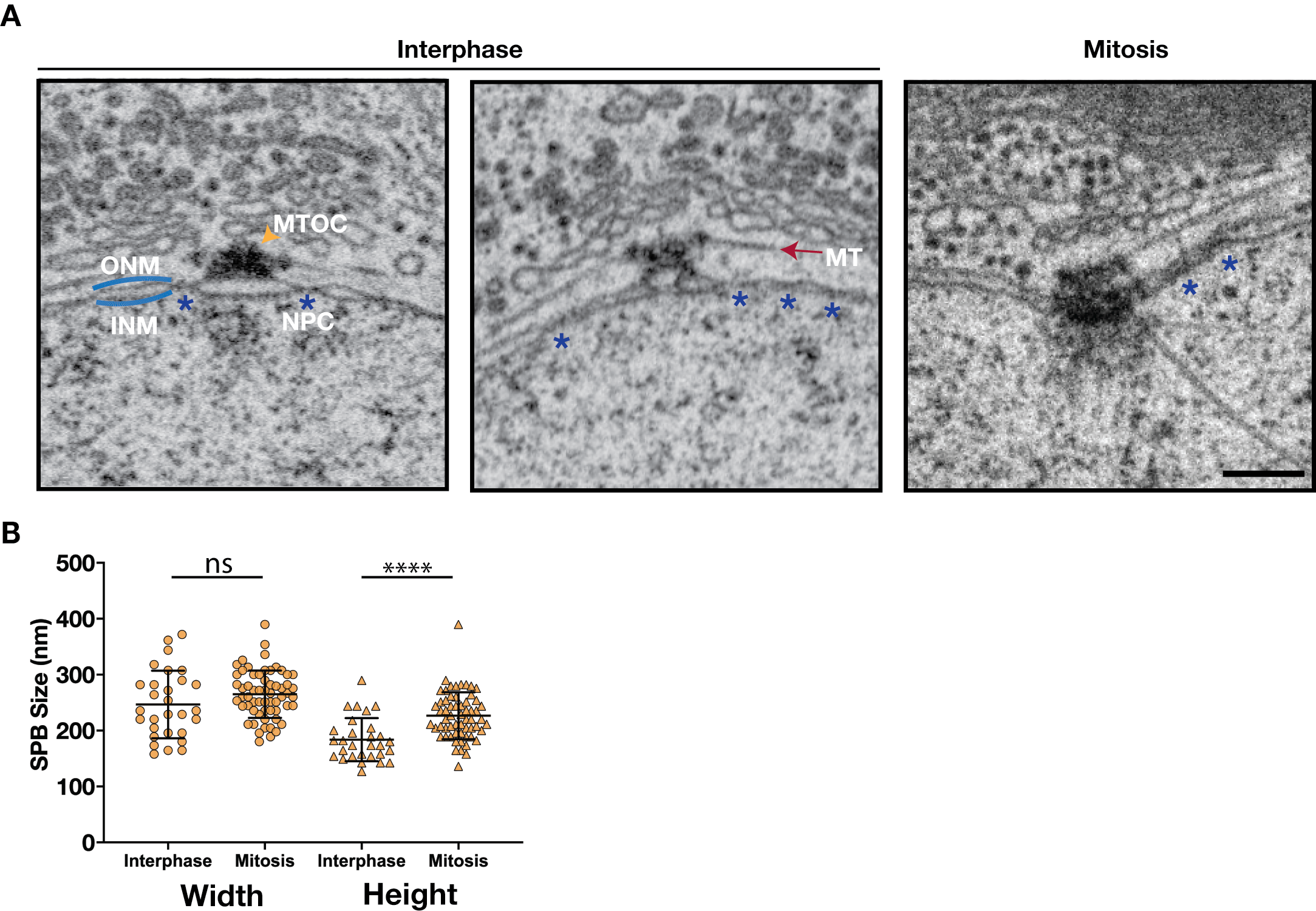
The MTOC of *S. arctica* is embedded in the nuclear envelope during mitosis. (**A**) FIB-SEM images showing different configurations of the S. *arctica* MTOC-NE interface, from outer nuclear membrane (ONM) association in interphase to inner nuclear membrane (INM) embedding in mitosis. Blue asterisks indicate nuclear pore complexes (NPCs). Scale bar = 300 nm. (**B**) Dot plot showing dimensions of *S. arctica*s MTOCs in interphase and mitosis. Width and height of MTOCs were measured from NHS-ester labelled U-ExM gels (n_interphase_ = 29, n_mitosis_ = 60) (ns-not significant; \*\*\*\**P* < 0.0001).

**Fig. S4.**
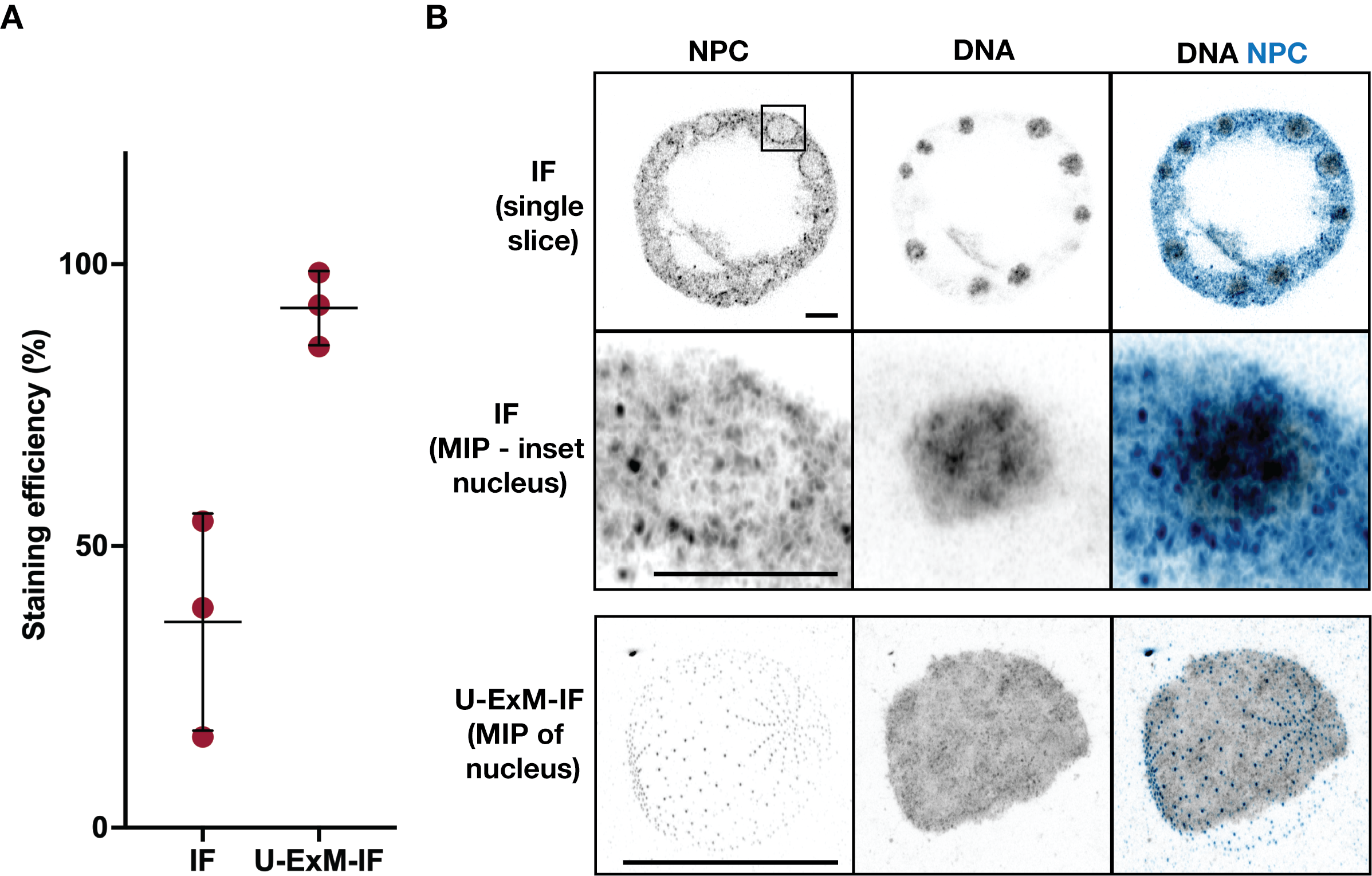
U-ExM improves immunostaining efficiency in Ichthyosporea. (**A**) Improvement in immunostaining efficiency when combined with U-ExM, measured as a percentage of cells labelled for tubulin (E7 antibody, DSHB), n = 3 experiments (n_IF_ = 199, 149, 164, n_UExM_ = 134, 82, 69). (**B**) U-ExM reveals ultrastructural details of the *S. arctica* nucleus in comparison with classical IF. Maximum intensity projections (MIP) of *S. arctica* nuclei stained either by IF (top) or IF post U-ExM (bottom) and labelled with MAb414 antibody (NPC, blue) and DNA (grey). Scale bar = 5 μm.

**Fig. S5.**
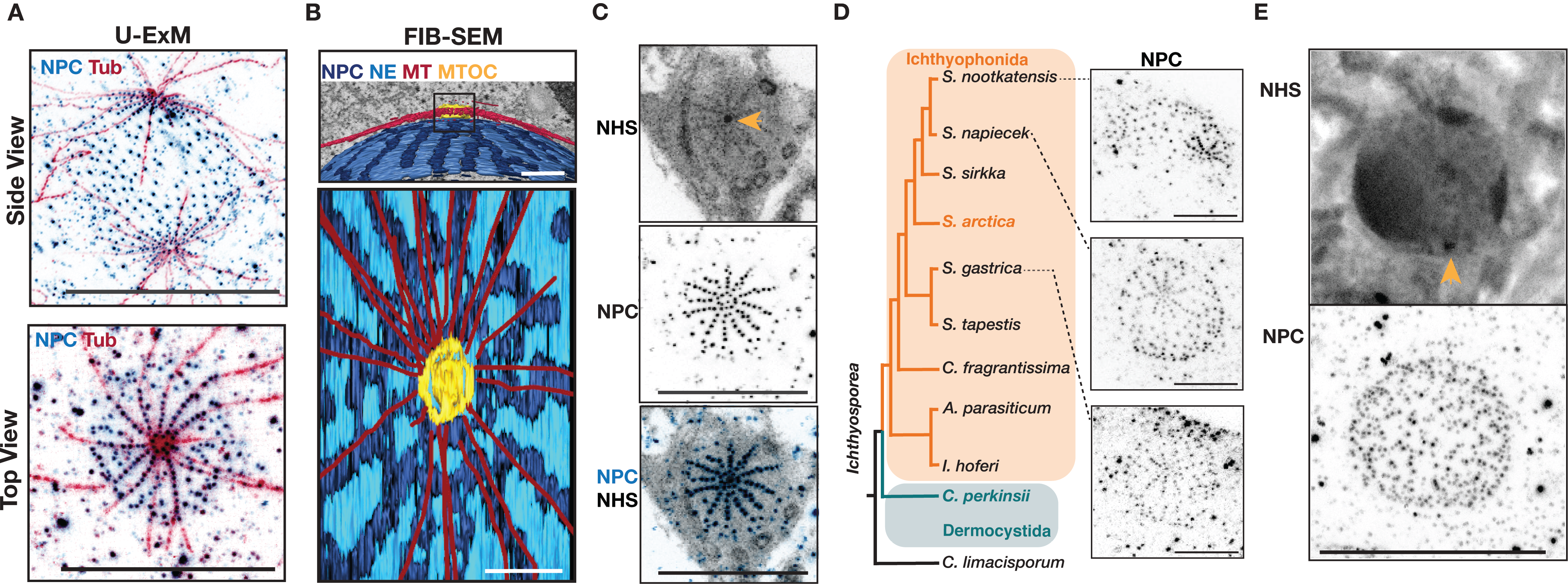
Radial distribution of NPCs surrounding the *S. arctica* MTOC. (**A**) Nuclear pore complexes (NPC-MAb414 antibody, blue) organised in radial arrays along MTs (tubulin, red). Images are maximum intensity projections of U-ExM stained *S. arctica* interphase nuclei. Scale bar = 5 μm. (**B**) 3D model of NPC arrays (dark blue) radiating from the MTOC (yellow) overlaid on an orthoslice of the *S. arctica* interphase nucleus imaged by FIB-SEM. Scale bar = 500 nm. (**C**) NPCs are present at the centre of the MTOC in a subpopulation of nuclei. Maximum intensity projections of U-ExM images labelled with pan protein label NHS ester (grey) and MAb414 antibody (NPC, blue). Yellow arrowhead indicates MTOC position. Scale bar = 5μm. (**D**) NPCs are organised in radial arrays across three distinct *Sphaeroforma* species. Images shown are maximum intensity projections of nuclei from *Sphaeroforma* sister species, *S. nootkatensis, S. napiecek* and *S. gastrica*. Scale bar = 2 μm. (**E**) NPCs are disorganised in *S. arctica* on prolonged microtubule inhibitor carbendazim (MBC) treatment (25 μg/ml MBC for 4 h). Images shown are maximum intensity projections of MBC-treated nuclei with pan protein labelling (NHS-ester) and MAb414 immunostaining. Yellow arrowhead indicates MTOC position. Scale bar = 5μm.

**Fig. S6.**
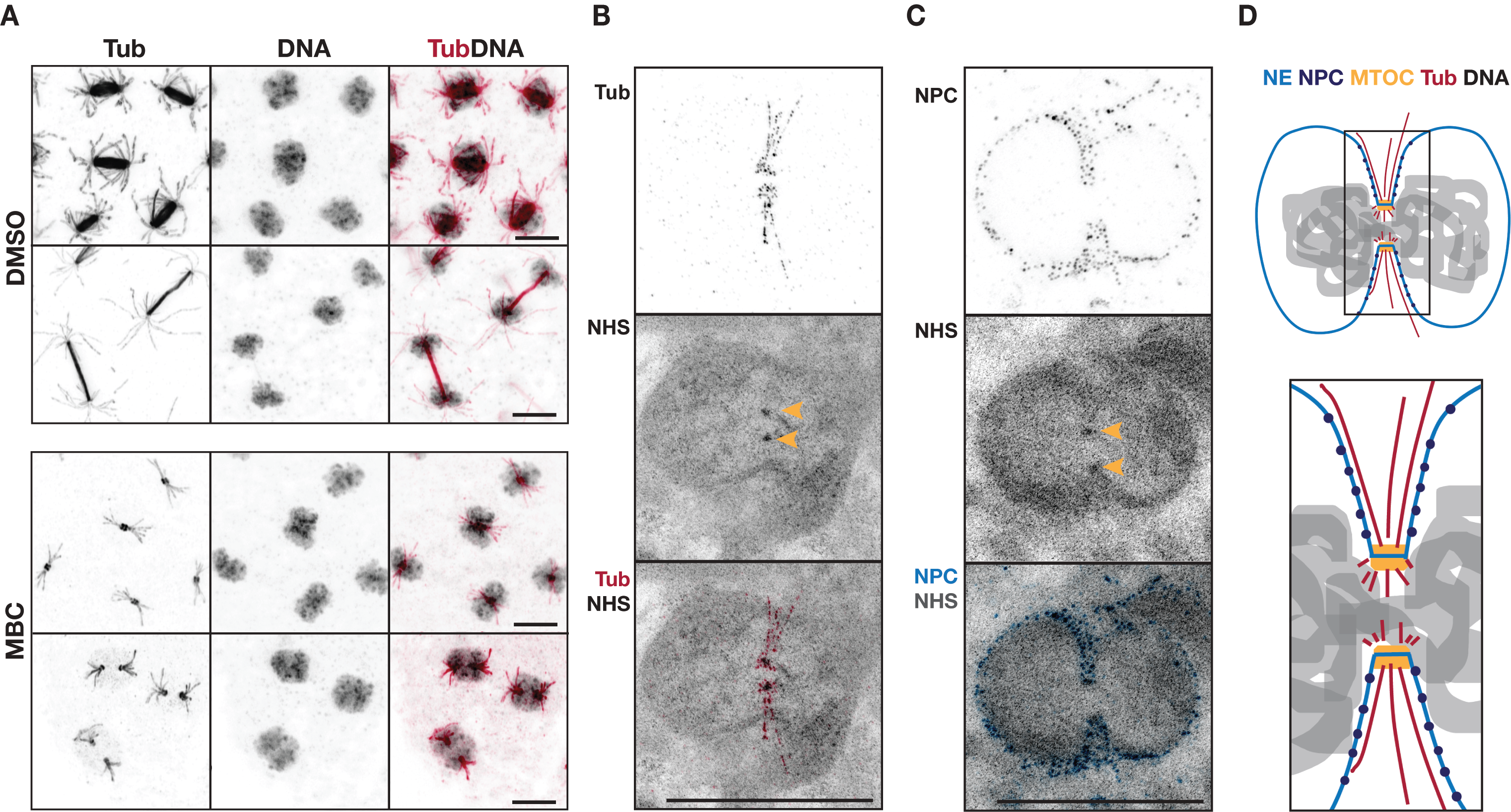
*S. arctica* nuclear remodelling in response to acute microtubule depolymerization. (**A**) *S. arctica* spindles collapse with acute microtubule inhibition (0.5 μg/ml MBC for 15 min). Maximum intensity projections of DMSO and MBC treated and immunolabeled *S. arctica* spindles at different stages of mitosis. Scale bar = 5 μm. Maximum intensity projections of U-ExM images of *S. arctica* nuclei with tubulin (Tub - red) and pan-labelling (NHS) shows loss of spindle MTs (**B**) while short astral MTs and (**C**) NPC radial arrays (blue) still persist. Scale bar = 2 μm. (**D**) Schematic representation of nuclear remodelling with increased polar indentation and spindle collapse in response to acute MT inhibitor treatment.

**Fig. S7.**
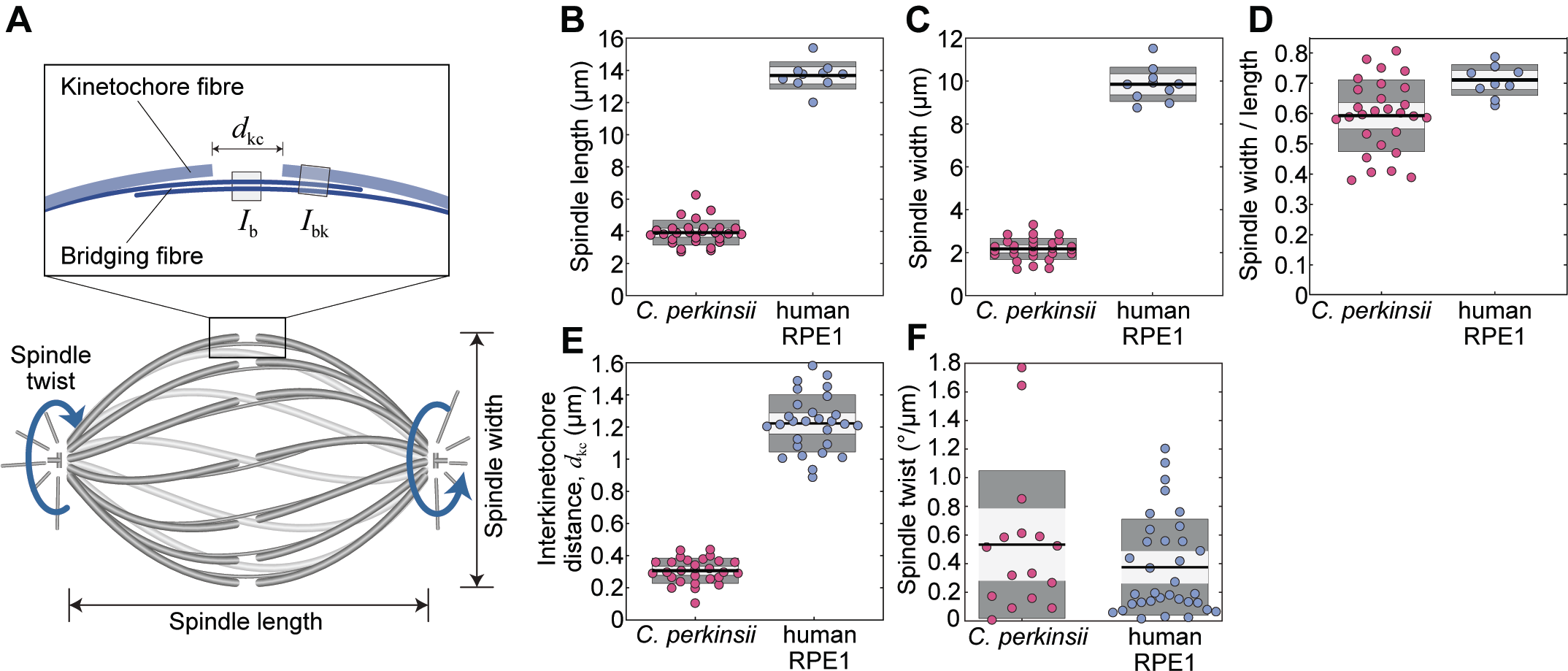
*C. perkinsii* shows features of human spindles. (**A**) Schematic showing features of mitotic spindles including kinetochore and bridging fibres, interkinetochore distance (d_kc_), twist, spindle length and width. Plot of (**B**) spindle length, (**C**) spindle width and (**D**) spindle width/length ratio, (**E**) interkinetochore distance and (**F**) absolute value of spindle twist in *C. perkinsii* and human RPE1 cells. Data for RPE1 cells in (**D**) and (**E**) was taken from Štimac et al., 2022 (*32, 69*).

**Fig. S8.**
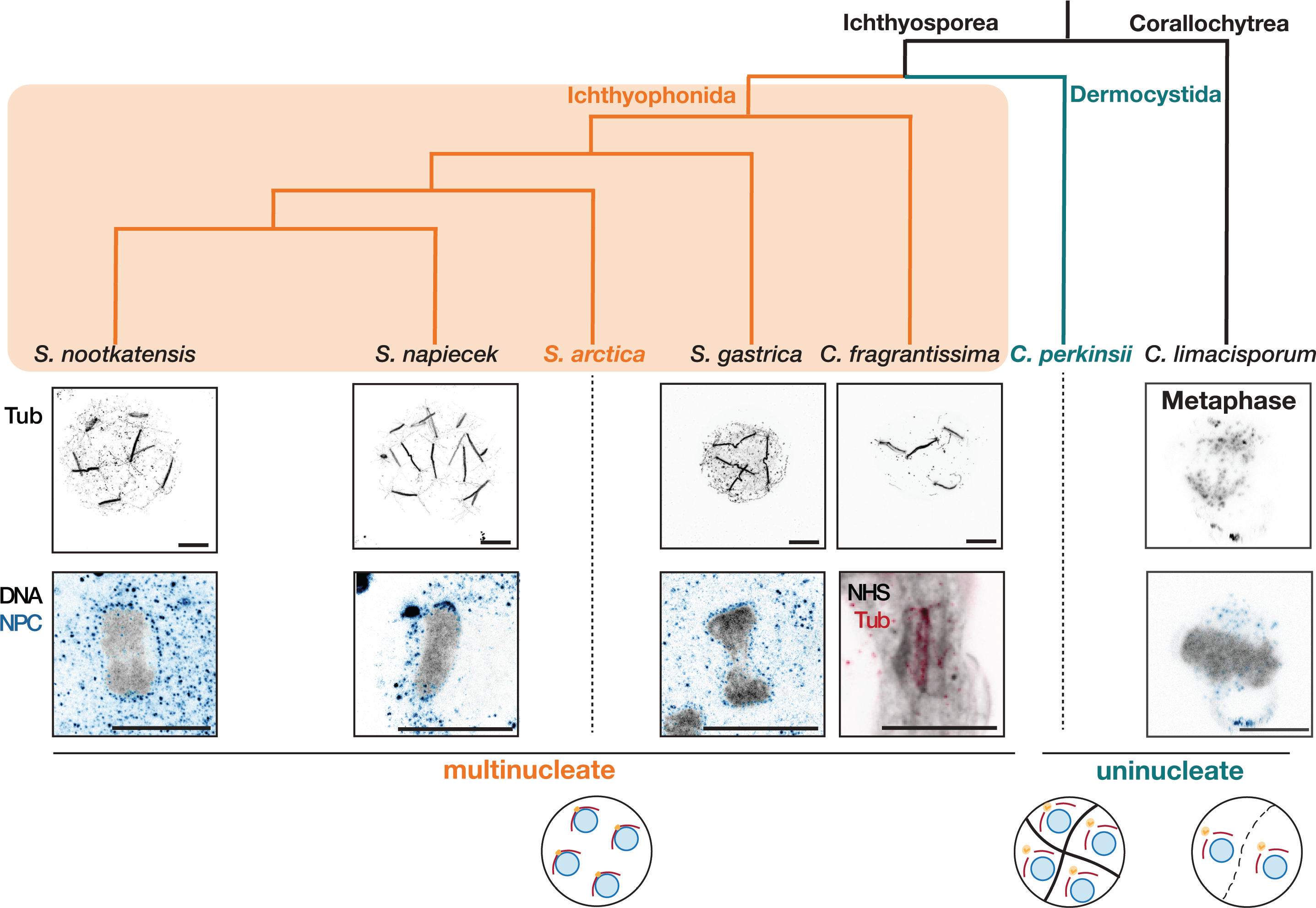
Mitotic strategies in Ichthyophonida, Dermocystida and Corallochytrea. Phylogenetic relationship of Ichthyophonida and Dermocystida within the Ichthyosporea. Ichthyophonida undergo closed mitosis. Central panel shows maximum intensity projections of U-ExM images of anaphase spindle architecture (top, Tubulin) and NPC organisation (bottom, NPC (blue) and DNA (grey)) in cells of representative Ichthyophonida and Corallochytrea species. Bottom panel shows the uni- or multinucleate life cycle of the selected species. Scale bar = 5 μm. Loss of NPCs through mitosis in *C. limacisporum* as seen in maximum intensity projections of *C. limacisporum* nuclei at prophase and metaphase labelled for tubulin and DNA along with MAb414 (NPC, blue) post U-ExM. Scale bar = 2 μm.

**Table S1:** Features of mitotic spindles in *C. perkinsii* and Human RPE-1 cells.

| *C. perkinsii* |  |  |  |  |  |  | Human RPE-1 |  |  |  |  |  |  |
| --- | --- | --- | --- | --- | --- | --- | --- | --- | --- | --- | --- | --- | --- |
| Lengt<br>h<br>(μm) | Widt<br>h<br>(μm) | Widt<br>h<br>/<br>lengt<br>h | dkc<br>(μm) | dkc /<br>spin<br>dle<br>lengt<br>h | Ib/<br>Ibk | Absol<br>ute<br>twist<br>($^{\circ}$/μm<br>) | Lengt<br>h<br>(μm) | Widt<br>h<br>(μm) | Widt<br>h<br>/<br>lengt<br>h | dkc<br>(μm) | dkc /<br>spin<br>dle<br>lengt<br>h | Ib/<br>Ibk | Absol<br>ute<br>twist<br>($^{\circ}$/μm<br>) |
| 3.91 ±<br>0.15<br>(28) | 2.28<br>±<br>0.09<br>(28) | 0.59<br>±<br>0.02<br>(28) | 0.31<br>±<br>0.01<br>(28) | 0.08<br>±<br>0.00<br>6<br>(28) | 0.41<br>±<br>0.03<br>(28) | 0.53 ±<br>0.13<br>(16) | 13.69<br>± 0.27<br>(10) | 9.73<br>±<br>0.25<br>(10) | 0.71<br>±<br>0.02<br>(10) | 1.22<br>±<br>0.03<br>(28) | 0.09<br>±<br>0.00<br>4<br>(10) | 0.22<br>±<br>0.03<br>(30) | 0.37 ±<br>0.06<br>(34) |

**Movie S1**: FIB-SEM volume of an interphase *S. arctica* nucleus.

**Movie S2**: Light-sheet imaging of *S. arctica* mitosis. sss

**Movie S3**: 3D reconstruction of *S. arctica* metaphase spindle shown in Fig. 2C. Tubulin (red), DNA (grey) and NPCs (blue).

**Movie S4**: 3D reconstruction of *S. arctica* anaphase spindle shown in Fig. 2C. Tubulin (red), DNA (grey) and NPCs (blue).

**Movie S5**: Serial tomogram of *S. arctica* mitotic nucleus.

**Movie S6**: Tomogram of *S. arctica* anaphase nucleus.

**Movie S7**: 3D reconstruction of *C. perkinsii* metaphase spindle shown in Fig. 3B. Tubulin (red), DNA (grey) and NPCs (blue).

**Movie S8**: 3D reconstruction of *C. perkinsii* anaphase spindle shown in Fig. 3B. Tubulin (red), DNA (grey) and NPCs (blue).

**Movie S9**: Serial tomogram of *C. perkinsii* mitotic nucleus.

**Data S1**: Eukprot IDs of homologs of MTOC, NE, MT and KT associated proteins in close relatives of animals.

## Notes

https://doi.org/10.6084/m9.figshare.c.6639812.v1

### Competing Interest Statement

The authors have declared no competing interest.

## References

1. G. Dey, B. Baum, Nuclear envelope remodelling during mitosis. Curr Opin Cell Biol. 70, 67–74 (2021).

2. M. Makarova, S. Oliferenko, Mixing and matching nuclear envelope remodeling and spindle assembly strategies in the evolution of mitosis. Current Opinion in Cell Biology. 41, 43–50 (2016).

3. M. W. Hetzer, T. C. Walther, I. W. Mattaj, PUSHING THE ENVELOPE: Structure, Function, and Dynamics of the Nuclear Periphery. Annu. Rev. Cell Dev. Biol. 21, 347–380 (2005).

4. M. Winey, D. Yarar, T. H. Giddings, D. N. Mastronarde, Nuclear Pore Complex Number and Distribution throughout the *Saccharomyces cerevisiae* Cell Cycle by Three-Dimensional Reconstruction from Electron Micrographs of Nuclear Envelopes. MBoC. 8, 2119–2132 (1997).

5. K. Tanaka, T. Kanbe, Mitosis in the fission yeast Schizosaccharomyces pombe as revealed by freeze-substitution electron microscopy. Journal of Cell Science. 80, 253–268 (1986).

6. G. Dey, S. Culley, S. Curran, U. Schmidt, R. Henriques, W. Kukulski, B. Baum, Closed mitosis requires local disassembly of the nuclear envelope. Nature. 585, 119–123 (2020).

7. C. P. C. De Souza, S. A. Osmani, Mitosis, Not Just Open or Closed. Eukaryotic Cell. 6, 1521–1527 (2007).

8. S. Sazer, M. Lynch, D. Needleman, Deciphering the Evolutionary History of Open and Closed Mitosis. Current Biology. 24, R1099–R1103 (2014).

9. H. Drechsler, A. D. McAinsh, Exotic mitotic mechanisms. Open Biol. 2, 120140 (2012).

10. I. B. Heath, “Variant Mitoses in Lower Eukaryotes: Indicators of the Evolution of Mitosis?” in International Review of Cytology (Elsevier, 1980; https://linkinghub.elsevier.com/retrieve/pii/S0074769608602351), vol. 64, pp. 1–80.

11. J. Wu, A. Akhmanova, Microtubule-Organizing Centers. Annu. Rev. Cell Dev. Biol. 33, 51–75 (2017).

12. G. J. P. L. Kops, B. Snel, E. C. Tromer, Evolutionary Dynamics of the Spindle Assembly Checkpoint in Eukaryotes. Current Biology. 30, R589–R602 (2020).

13. J. J. Vicente, L. Wordeman, Mitosis, microtubule dynamics and the evolution of kinesins. Exp Cell Res. 334, 61–69 (2015).

14. P. De Magistris, W. Antonin, The Dynamic Nature of the Nuclear Envelope. Current Biology. 28, R487–R497 (2018).

15. R. Ungricht, U. Kutay, Mechanisms and functions of nuclear envelope remodelling. Nat Rev Mol Cell Biol. 18, 229–245 (2017).

16. J. J. Hooff, E. Tromer, L. M. Wijk, B. Snel, G. J. Kops, Evolutionary dynamics of the kinetochore network in eukaryotes as revealed by comparative genomics. EMBO Rep. 18, 1559–1571 (2017).

17. E. C. Tromer, J. J. E. van Hooff, G. J. P. L. Kops, B. Snel, Mosaic origin of the eukaryotic kinetochore. Proc Natl Acad Sci USA. 116, 12873–12882 (2019).

18. I. A. Drinnenberg, B. Akiyoshi, “Evolutionary Lessons from Species with Unique Kinetochores” in Centromeres and Kinetochores, B. E. Black, Ed. (Springer International Publishing, Cham, 2017; http://link.springer.com/10.1007/978-3-319-58592-5_5), vol. 56 of *Progress in Molecular and Subcellular Biology*, pp. 111–138.

19. D. Rüthnick, E. Schiebel, Duplication and Nuclear Envelope Insertion of the Yeast Microtubule Organizing Centre, the Spindle Pole Body. Cells. 7, 42 (2018).

20. S. Otsuka, A. M. Steyer, M. Schorb, J.-K. Hériché, M. J. Hossain, S. Sethi, M. Kueblbeck, Y. Schwab, M. Beck, J. Ellenberg, Postmitotic nuclear pore assembly proceeds by radial dilation of small membrane openings. Nat Struct Mol Biol. 25, 21–28 (2018).

21. A. Sebé-Pedrós, B. M. Degnan, I. Ruiz-Trillo, The origin of Metazoa: a unicellular perspective. Nat Rev Genet. 18, 498–512 (2017).

22. M. A. Naranjo-Ortiz, T. Gabaldón, Fungal evolution: diversity, taxonomy and phylogeny of the Fungi. Biol Rev Camb Philos Soc. 94, 2101–2137 (2019).

23. G. Torruella, A. de Mendoza, X. Grau-Bové, M. Antó, M. A. Chaplin, J. del Campo, L. Eme, G. Pérez-Cordón, C. M. Whipps, K. M. Nichols, R. Paley, A. J. Roger, A. Sitjà-Bobadilla, S. Donachie, I. Ruiz-Trillo, Phylogenomics Reveals Convergent Evolution of Lifestyles in Close Relatives of Animals and Fungi. Current Biology. 25, 2404–2410 (2015).

24. T. Brunet, N. King, The Origin of Animal Multicellularity and Cell Differentiation. Developmental Cell. 43, 124–140 (2017).

25. O. Dudin, A. Ondracka, X. Grau-Bové, A. A. Haraldsen, A. Toyoda, H. Suga, J. Bråte, I. Ruiz-Trillo, A unicellular relative of animals generates a layer of polarized cells by actomyosin-dependent cellularization. eLife. 8, e49801 (2019).

26. B. McCartney, O. Dudin, Cellularization across eukaryotes: Conserved mechanisms and novel strategies. Current Opinion in Cell Biology. 80, 102157 (2023).

27. K. D. Arkush, L. Mendoza, M. A. Adkison, R. P. Hedrick, Observations on the Life Stages of Sphaerothecum destruens n. g., n. sp., a Mesomycetozoean Fish Pathogen Formally Referred to as the Rosette Agent. J Eukaryotic Microbiology. 50, 430–438 (2003).

28. L. Mendoza, J. W. Taylor, L. Ajello, The Class Mesomycetozoea: A Heterogeneous Group of Microorganisms at the Animal-Fungal Boundary. Annu. Rev. Microbiol. 56, 315–344 (2002).

29. C. Schilde, P. Schaap, The Amoebozoa. Methods Mol Biol. 983, 1–15 (2013).

30. J. J. van Hooff, E. Tromer, L. M. van Wijk, B. Snel, G. J. Kops, Evolutionary dynamics of the kinetochore network in eukaryotes as revealed by comparative genomics. EMBO Rep. 18, 1559–1571 (2017).

31. L. E. van Rooijen, E. C. Tromer, J. J. E. van Hooff, G. J. P. L. Kops, B. Snel, Increased Sampling and Intracomplex Homologies Favor Vertical Over Horizontal Inheritance of the Dam1 Complex. Genome Biol Evol. 15, evad017 (2023).

32. V. Štimac, I. Koprivec, M. Manenica, J. Simunić, I. M. Tolić, Augmin prevents merotelic attachments by promoting proper arrangement of bridging and kinetochore fibers. Elife. 11, e83287 (2022).

33. Z. She, Y. Wei, Y. Lin, Y. Li, M. Lu, Mechanisms of the Ase1/PRC1/MAP65 family in central spindle assembly. Biol Rev. 94, 2033–2048 (2019).

34. I. Tikhonenko, D. K. Nag, D. N. Robinson, M. P. Koonce, Microtubule-Nucleus Interactions in Dictyostelium discoideum Mediated by Central Motor Kinesins. Eukaryotic Cell. 8, 723–731 (2009).

35. A. Ondracka, O. Dudin, I. Ruiz-Trillo, Decoupling of Nuclear Division Cycles and Cell Size during the Coenocytic Growth of the Ichthyosporean Sphaeroforma arctica. Curr Biol. 28, 1964–1969.e2 (2018).

36. M. Olivetta, O. Dudin, The nuclear-to-cytoplasmic ratio drives cellularization in the close animal relative Sphaeroforma arctica. Current Biology, S096098222300307X (2023).

37. A. Gupta, D. Kitagawa, Ultrastructural diversity between centrioles of eukaryotes. The Journal of Biochemistry. 164, 1–8 (2018).

38. W. L. Marshall, M. L. Berbee, Comparative Morphology and Genealogical Delimitation of Cryptic Species of Sympatric Isolates of Sphaeroforma (Ichthyosporea, Opisthokonta). Protist. 164, 287–311 (2013).

39. X. Grau-Bové, G. Torruella, S. Donachie, H. Suga, G. Leonard, T. A. Richards, I. Ruiz-Trillo, Dynamics of genomic innovation in the unicellular ancestry of animals. eLife. 6, e26036 (2017).

40. A. Kożyczkowska, S. R. Najle, E. Ocaña-Pallarès, C. Aresté, V. Shabardina, P. S. Ara, I. Ruiz-Trillo, E. Casacuberta, Stable transfection in protist Corallochytrium limacisporum identifies novel cellular features among unicellular animals relatives. Curr Biol. 31, 4104–4110.e5 (2021).

41. R. A. Coss, Mitosis in Chlamydomonas reinhardtii basal bodies and the mitotic apparatus. J Cell Biol. 63, 325–329 (1974).

42. R. Rashpa, M. Brochet, “Ultrastructure expansion microscopy of *Plasmodium* gametocytes reveals the molecular architecture of a microtubule organisation centre coordinating mitosis with axoneme assembly (preprint, Microbiology, 2021), doi:10.1101/2021.07.21.453039.

43. M. J. Powell, MITOSIS IN THE AQUATIC FUNGUS RHIZOPHYDIUM SPHEROTHECA (CHYTRIDIALES). American Journal of Botany. 67, 839–853 (1980).

44. J. Boruc, X. Zhou, I. Meier, Dynamics of the Plant Nuclear Envelope and Nuclear Pore. Plant Physiol. 158, 78–86 (2012).

45. R. C. Brown, B. E. Lemmon, The Pleiomorphic Plant MTOC: An Evolutionary Perspective. Journal of Integrative Plant Biology. 49, 1142–1153 (2007).

46. K. R. Katsani, R. E. Karess, N. Dostatni, V. Doye, In Vivo Dynamics of *Drosophila* Nuclear Envelope Components. MBoC. 19, 3652–3666 (2008).

47. C. Roubinet, B. Decelle, G. Chicanne, J. F. Dorn, B. Payrastre, F. Payre, S. Carreno, Molecular networks linked by Moesin drive remodeling of the cell cortex during mitosis. J Cell Biol. 195, 99–112 (2011).

48. T. Duan, R. Cupp, P. K. Geyer, Drosophila female germline stem cells undergo mitosis without nuclear breakdown. Current Biology. 31, 1450–1462.e3 (2021).

49. A. R. Gerhold, J.-C. Labbé, R. Singh, Uncoupling cell division and cytokinesis during germline development in metazoans. Front. Cell Dev. Biol. 10, 1001689 (2022).

50. H. S. Seidel, T. A. Smith, J. K. Evans, J. Q. Stamper, T. G. Mast, J. Kimble, C. elegans germ cells divide and differentiate in a folded tissue. Developmental Biology. 442, 173–187 (2018).

51. C. Lang, S. Grava, T. van den Hoorn, R. Trimble, P. Philippsen, S. L. Jaspersen, Mobility, Microtubule Nucleation and Structure of Microtubule-organizing Centers in Multinucleated Hyphae of *Ashbya gossypii*. MBoC. 21, 18–28 (2010).

52. L. Solnica-Krezel, T. G. Burland, W. F. Dove, Variable pathways for developmental changes of mitosis and cytokinesis in Physarum polycephalum. J Cell Biol. 113, 591–604 (1991).

53. T. G. Burland, L. Solnica, W. F. Dove, Patterns of Inheritance, Development and the Mitotic Cycle in the Protist Physamm polycephalum, 69.

54. K. Mitic, M. Grafe, P. Batsios, I. Meyer, Partial Disassembly of the Nuclear Pore Complex Proteins during Semi-Closed Mitosis in Dictyostelium discoideum. Cells. 11, 407 (2022).

55. Y. Gu, S. Oliferenko, Comparative biology of cell division in the fission yeast clade. Curr Opin Microbiol. 28, 18–25 (2015).

56. U. Theisen, A. Straube, G. Steinberg, Dynamic rearrangement of nucleoporins during fungal “open” mitosis. Mol Biol Cell. 19, 1230–1240 (2008).

57. D. J. Richter, C. Berney, J. F. H. Strassert, Y.-P. Poh, E. K. Herman, S. A. Muñoz-Gómez, J. G. Wideman, F. Burki, C. De Vargas, EukProt: A database of genome-scale predicted proteins across the diversity of eukaryotes. Peer Community Journal. 2, e56 (2022).

58. S. R. Eddy, Accelerated Profile HMM Searches. PLoS Comput Biol. 7, e1002195 (2011).

59. A. Bateman, L. Coin, R. Durbin, R. D. Finn, V. Hollich, S. Griffiths-Jones, A. Khanna, M. Marshall, S. Moxon, E. L. L. Sonnhammer, D. J. Studholme, C. Yeats, S. R. Eddy, The Pfam protein families database. Nucleic Acids Res. 32, D138–141 (2004).

60. K. Katoh, D. M. Standley, MAFFT Multiple Sequence Alignment Software Version 7: Improvements in Performance and Usability. Molecular Biology and Evolution. 30, 772–780 (2013).

61. S. Capella-Gutiérrez, J. M. Silla-Martínez, T. Gabaldón, trimAl: a tool for automated alignment trimming in large-scale phylogenetic analyses. Bioinformatics. 25, 1972–1973 (2009).

62. L.-T. Nguyen, H. A. Schmidt, A. Von Haeseler, B. Q. Minh, IQ-TREE: A Fast and Effective Stochastic Algorithm for Estimating Maximum-Likelihood Phylogenies. Molecular Biology and Evolution. 32, 268–274 (2015).

63. S. Kalyaanamoorthy, B. Q. Minh, T. K. F. Wong, A. Von Haeseler, L. S. Jermiin, ModelFinder: fast model selection for accurate phylogenetic estimates. Nat Methods. 14, 587–589 (2017).

64. D. T. Hoang, O. Chernomor, A. Von Haeseler, B. Q. Minh, L. S. Vinh, UFBoot2: Improving the Ultrafast Bootstrap Approximation. Molecular Biology and Evolution. 35, 518–522 (2018).

65. D. Gambarotto, F. U. Zwettler, M. Le Guennec, M. Schmidt-Cernohorska, D. Fortun, S. Borgers, J. Heine, J.-G. Schloetel, M. Reuss, M. Unser, E. S. Boyden, M. Sauer, V. Hamel, P. Guichard, Imaging cellular ultrastructures using expansion microscopy (U-ExM). Nat Methods. 16, 71–74 (2019).

66. J. Schindelin, I. Arganda-Carreras, E. Frise, V. Kaynig, M. Longair, T. Pietzsch, S. Preibisch, C. Rueden, S. Saalfeld, B. Schmid, J.-Y. Tinevez, D. J. White, V. Hartenstein, K. Eliceiri, P. Tomancak, A. Cardona, Fiji: an open-source platform for biological-image analysis. Nat Methods. 9, 676–682 (2012).

67. C. T. Rueden, J. Schindelin, M. C. Hiner, B. E. DeZonia, A. E. Walter, E. T. Arena, K. W. Eliceiri, ImageJ2: ImageJ for the next generation of scientific image data. BMC Bioinformatics. 18, 529 (2017).

68. M. Novak, B. Polak, J. Simunić, Z. Boban, B. Kuzmić, A. W. Thomae, I. M. Tolić, N. Pavin, The mitotic spindle is chiral due to torques within microtubule bundles. Nat Commun. 9, 3571 (2018).

69. M. Trupinić, B. Kokanović, I. Ponjavić, I. Barišić, S. Šegvić, A. Ivec, I. M. Tolić, The chirality of the mitotic spindle provides a mechanical response to forces and depends on microtubule motors and augmin. Curr Biol. 32, 2480–2493.e6 (2022).

70. J. Kajtez, A. Solomatina, M. Novak, B. Polak, K. Vukušić, J. Rüdiger, G. Cojoc, A. Milas, I. Šumanovac Šestak, P. Risteski, F. Tavano, A. H. Klemm, E. Roscioli, J. Welburn, D. Cimini, M. Glunčić, N. Pavin, I. M. Tolić, Overlap microtubules link sister k-fibres and balance the forces on bi-oriented kinetochores. Nat Commun. 7, 10298 (2016).

71. B. Polak, P. Risteski, S. Lesjak, I. M. Tolić, PRC1-labeled microtubule bundles and kinetochore pairs show one-to-one association in metaphase. EMBO Rep. 18, 217–230 (2017).

72. M. W. Elting, M. Prakash, D. B. Udy, S. Dumont, Mapping Load-Bearing in the Mammalian Spindle Reveals Local Kinetochore Fiber Anchorage that Provides Mechanical Isolation and Redundancy. Curr Biol. 27, 2112–2122.e5 (2017).

73. S. Suresh, S. Markossian, A. H. Osmani, S. A. Osmani, Mitotic nuclear pore complex segregation involves Nup2 in Aspergillus nidulans. Journal of Cell Biology. 216, 2813–2826 (2017).

74. J. Matković, S. Ghosh, M. Ćosić, N. Pavin, I. M. Tolić, “Kinetochore- and chromosome-driven transition of microtubules into bundles promotes spindle assembly” (preprint, Cell Biology, 2022), doi:10.1101/2022.02.25.481924.

75. M. Jagrić, P. Risteski, J. Martinčić, A. Milas, I. M. Tolić, Optogenetic control of PRC1 reveals its role in chromosome alignment on the spindle by overlap length-dependent forces. Elife. 10, e61170 (2021).

76. P. Risteski, D. Božan, M. Jagrić, A. Bosilj, N. Pavin, I. M. Tolić, Length-dependent poleward flux of sister kinetochore fibers promotes chromosome alignment. Cell Rep. 40, 111169 (2022).

77. K. Vukušić, R. Buđa, A. Bosilj, A. Milas, N. Pavin, I. M. Tolić, Microtubule Sliding within the Bridging Fiber Pushes Kinetochore Fibers Apart to Segregate Chromosomes. Dev Cell. 43, 11–23.e6 (2017).

78. Z. Jiang, S. Zhang, Y. M. Lee, X. Teng, Q. Yang, Y. Toyama, Y.-C. Liou, Hyaluronan-Mediated Motility Receptor Governs Chromosome Segregation by Regulating Microtubules Sliding Within the Bridging Fiber. Adv Biol (Weinh). 5, e2000493 (2021).

